# Computational vaccinology approach: Designing an efficient multi-epitope peptide vaccine against *Cryptococcus neoformans var. grubii’s* heat shock 70KDa protein

**DOI:** 10.1101/534008

**Authors:** Reham M. Elhassan, Nagla M. Alsony, Khadeejah M. Othman, Duaa T. Izz-Aldin, Tamadour A. Alhaj, Abdelrahman A. Ali, Lena A. Abashir, Omar H. Ahmed, Mohammed A. Hassan

## Abstract

**Introduction:** Cryptococcosis is a ubiquitous opportunistic fungal disease caused by Cryptococcus neoformans var. grubii. It has high global morbidity and mortality among HIV patients and none-HIV carriers with 99% and 95% respectively. Furthermore, the increasing prevalence of undesired toxicity profile of antifungal, multi-drug resistant organism, and the scarcity of FDA authorized vaccines, where the hallmark in the present days. This study was undertaken to design a reliable multi-epitope peptide vaccine against highly conserved immunodominant heat shock 70KDa protein of Cryptococcus neoformans var. grubii that covers a considerable digit of the world population through implementing computational vaccinology approach.

**Materials and Methods:** A total of 38 Sequences of Cryptococcus neoformans var. grubii’s heat shock 70KDa protein were retrieved from NCBI protein database. Different prediction tools were used to analyze the aforementioned protein at Immune Epitope Database (IEDB) to discriminate the most promising T-cell and B-cell epitopes. Then the proposed epitopes were subjected to Population coverage analysis tool to compute global population’s coverage. Finally, the projected epitopes were ranked based on their scores and binding modes through using Moe 2007 program.

**Outstanding Results and Conclusion:** Our prime vaccine candidate was a putative ten promising epitopes (ANYVQASEK, NYVQASEK, KSVEKPAS, TPQQPPAQ, YVYDTRGKL, FYRQGAFEL, FTQLVAAYL, FFGGKVLNF, FDYALVQHF, and FINAQLVDV). Together, these epitopes are forecasted to trigger T lymphocytes, B lymphocytes, and immunological memory with overall population coverage above 90%. Accordingly, our in silico vaccine is expected to be the future multi-epitope peptide vaccine against Cryptococcus neoformans var. grubii’s heat shock 70KDa protein that covers a significant figure of the entire world citizens. Therefore, there is a definite need for experimental validation for the carefully chosen vaccine candidates in vitro and in vivo to fortify their antigenic and immunogenic potentials. Additionally, further computational studies are needed to be conducted in pathogens-derived Heat shock 70KDa protein family, as it believed to find universal epitopes that might be overlapped with other pathogens-derived Hsp70.

## 1. Introduction

Cryptococcosis is a ubiquitous opportunistic infection caused by *Cryptococcus neoformans* that causes life-threatening pneumonia and meningoencephalitis in immunocompromised patients (1–41). Cryptococcosis is considered as one of the most predominant causes of death globally with estimated annual mortality of 624,700 cases (5, 27, 35, 39, 42–47). Commonly *Cryptococcus neoformans var. grubii* strains (serotype A) are more virulent, widely distributed all over the world, and cause 99% of infections in HIV patients and 95% of the none-HIV carriers (5, 25, 27, 44, 48–57).

Clinical and experimental evidence suggests that cell-mediated immunity play a crucial contribution to host defense against intracellular cryptococcosis, because of its extended immunological response half-life, and the antigen can easily escape the secreted antibodies (3, 4, 31, 58–61). Nevertheless, this does not contradict the importance of B-cell response, which is considered as a vital mechanism for inducing protection against cryptococcosis in individuals with impaired cell-mediated immunity (62–65), When it comes to virulence factors, given that *C. neoformans* expresses a significant number of virulence agents that could help the parasite to evade and evoke host immunity (7, 46, 66, 67). Interestingly, Heat shock 70KDa protein (Hsp70) found to be one of the novel immunogenic proteins that trigger a cellular and humoral response against all *C. neoformans var. grubii* strains (7, 46, 66, 68–76). Throughout evolution, the aforementioed protein is highly conserved cell-surface protein, and widely expressed in *Plasmodium, Trypanosoma, Schistosoma, Leishmania, aspergilla, candidia, histoplasma*, and *Mycobacterium* species. Thereby, Hsps70 are thought to be a universal target with other pathogens-derived Hsp70 (77–89).

Multi-epitope peptide vaccines are immune-proteome derived–pathogen, compose of several peptide fragments of T-cell and B-cell epitopes that able to evoke highly targeted lymphocytes response and immunological memory (90–100). On one hand, peptide-based vaccines potentially have several benefits include they are cost-effective and easy in manufacturing process, highly stable, and diminished toxicity are generally confirmed in clinical studies. In addition, peptide vaccines have no concerns in targeting diseases from virus infection to Alzheimer disease or allergy, and most vitally allow the optimization for a specific population; permitting the avoidance of adverse conventional vaccine reactions (58, 59, 91, 95, 101–108).

On the other hand, the conventional vaccines are not practical for use in immunocompromised patients because of its drawbacks such as infection and death are the major concerns, overall low levels of immunogenicity, and may evoke hypersensitivity reactions (91, 106).

Cryptococcal infections have become more prevalent in recent years due to the increase number of immunocompromised patients with AIDS (62). Current antifungal therapies are not effective in eradicating the pathogens because of the precipitation of a significant toxicity as in life-threatening Cryptococcus-related immune reconstitution inflammatory syndrome, which is detected mostly in HIV individuals whom are on antiviral with antifungal therapy (23, 32, 109–117). In addition, the widespread use of intravascular catheters and irrational use of broad spectrum antibiotic has led to the increasing incidence of multi-drug resistant organisms, serious opportunistic, and nosocomial infections, eventually may necessitate a prolonged therapy or experience disease relapses (32, 34, 62, 118–122). Yet, to date there is no FDA authorized vaccine available to combat cryptococcosis. (9, 22, 34, 76, 110–113, 118–121, 123). Taken together, what stated above where the central question in this study. Thus, this underlying an urgent need for designing a reliable immunome-derived epitope-driven vaccine using computational vaccinology approach through mapping the parasite’s antigenic determinants, which are the minimal regions of the antigen that would bind to specific receptors on lymphocytes or to secreted antibodies, thereby eliciting cellular and humoral immunity. (9, 59, 107, 108, 124–128).

In this study, we aim to design a multi-epitope peptide vaccine by implementing an emerging approach in computational vaccinology. An ideal vaccine of highly conserved immunodominant B and T lymphocyte epitopes with a wide global population coverage design against *Cryptococcus neoformans var. grubii*. Accordingly, this is the first study that provides an exciting opportunity to utilize heat shock 70KDa protein as an attractive immune-proteomic factor that able to stimulate desirable immune responses against cryptococcosis.

## 2. Methodology

The flow chart demonstrates the overall process of peptide vaccine designing is illustrated in Figure 1.

### 2.1 Protein sequence retrieval

A total of 38 heat shock 70KDa protein sequences with a length of 773 mer were retrieved in FASTA format from National Centre for Biotechnology Information (NCBI), protein database (Accession No.XP_012053205.1) on 30^th^ July 2018 at (https://www.ncbi.nlm.nih.gov/protein).

### 2.2 Determination of the conserved regions

The retrieved sequences were aligned to allocate the conserved regions using multiple sequence alignment (MSA). The retrieved antigen sequences were run against the NCBI Reference Protein (RefSeq) using ClustalW as implemented in BioEdit sequence alignment editor software Version 7.2.5 (129). After which, the epitope candidates were analyzed by a variety of prediction methods through Immune Epitope Database IEDB analysis resource tools (http://www.iedb.org/).

### 2.3 B-cell epitope prediction

B-cell epitopes are determinants on the surface of pathogens that interact with B cell receptors. B-cell epitopes can be either continuous or discontinuous. Approximately, 10% of B-cell epitopes are continuous, consisting of a linear stretch of amino acids along the polypeptide chain. Thus, the majority of B-cell epitopes are a discontinuous or conformational structure (124, 130–132).

#### 2.3.1 Continuous B-cell epitope prediction

Analysis of epitopes binding affinity to B-cell were assessed by the IEDB B-cell epitope prediction tool at (http://tools.iedb.org/bcell/) on 30^th^ July 2018. The classical propensity scale methods and hidden Markov model programmed analysis resource were applied from IEDB to fulfill the following physiochemical criteria (Linearity, Surface accessibility, and immunogenicity)

##### 2.3.1.1 Prediction of linear B-cell epitopes

BepiPred test was conducted as a linear B-cell epitopes prediction method to sort out the linear conserved regions with a default threshold value of 0.249 (133–135).

##### 2.3.1.2 Prediction of surface accessibility

Emini surface accessibility test was implemented as surface B-cell epitopes prediction method to discriminate the surface conserved epitopes with a default cut-off value of 1.000 (136).

##### 2.3.1.3 Prediction of antigenicity

The kolaskar and tongaonker antigenicity method was used to differentiate the immunogenic sites with a default cut-off value of 1.024 (137).

#### 2.3.2 Discontinuous B-cell epitope prediction

The reference sequence of heat shock 70KDa protein was subjected to Swiss model on 21^th^ October 2018 in order to get the 3D structure (138–142). Based on the geometrical properties of protein structure, the discontinuous B-cell epitopes were filtered out at ElliPro prediction tool after submission of the modeled 3D structure at (http://tools.iedb.org/ellipro/). ElliPro implements three algorithms performing the approximation of the protein shape as an ellipsoid, calculation of residue protrusion index (PI) and clustering of neighboring residues based on their PI values. The minimum score and maximum distance (Angstrom) were calibrated in the default mode with score of 0.5, and 6, respectively (143).

### 2.4 Prediction of T-cell epitopes

T-cells identify antigens as short peptide segment in association with MHC molecules on antigen-presenting cells. There are two categories of T-cells:

1. CD8+ T cytotoxic cells, which recognize peptides displayed by MHC-I molecules,
2. CD4+ T helper cells, which recognize epitopes in association with MHC-II molecules.

As opposed to B-cell epitopes, T-cell epitopes only recognize linear peptides. MHC-I binding predictions are now very strong and have wide allelic coverage by integration with predictions of proteasomal cleavage and TAP binding sites. MHC-II binding predictions are not as well developed as MHC-I binding predictions, yet it is still developing at a fast rate (130).

#### 2.4.1 Prediction of MHC-I binding profile for conserved epitopes

Analysis of epitopes binding to MHC-I molecules were assessed by the IEDB MHC-I prediction tool at (http://tools.iedb.org/mhci/) on 13^th^ September 2018. Artificial Neural Network (ANN) version 2.2 was chosen as Prediction method as it depends on the median inhibitory concentration (IC_50_) (144–149). Since the functional cleft of MHC-I molecules is closed and can only accommodate short peptides ranging from 9 to 11 amino acids; all epitopes lengths were set to the optimum length of 9mers (150, 151). As the predicted output is given in units of IC**_50_** nM, absolute binding affinity threshold correlates better with immunogenicity (152). Therefore, lower IC_50_ value indicates greater binding affinity and vice versa. As a rough protocol, all conserved epitopes with IC_50_ score less than 50 nM have high affinity, less than 500 nM intermediate affinity and less than 5000 nM low affinity (153). Thus, Conserved promiscuous epitopes at score equal or less than 500 IC**_50_** are selected for further analysis whereas epitopes with IC_50_ greater than 500 were omitted.

#### 2.4.2 Prediction of MHC-II binding profile for conserved epitopes

In terms of analysis of MHC-II epitope, the selected candidates were assessed by the IEDB MHC-II prediction tool at (http://tools.iedb.org/mhcii/) on 14^th^ September 2018. Unlike MHC-I, MHC-II has a flexible pocket, therefore accommodates peptides of varying lengths, typically 12 to 26 mer. Regarding binding groove in MHC class II molecules, there are series of polymorphic pockets and plateaus that interact with several side chains of the peptide core sequence, providing the specificity of the MHC-peptide interaction. The consensus sequence of the peptides is set to be 9 mer (154). For MHC-II binding affinity profile, the most frequent human allele references set were used. NN-algin was chosen as Prediction method as it depends on the median inhibitory concentration (IC_50_) (155). Finally, all Conserved Immunodominant peptides at score equal or less than 100 median inhibitory concentration (IC_50_) were selected for further mapping analysis whereas epitopes with IC_50_ greater than 100 were dismissed (156).

### 2.5 Population coverage calculation

To ensure the universal coverage within a heterogeneous populations, it is crucial to calculate global population coverage for the chosen epitopes since the HLAs are among the most polymorphic proteins and varies among different geographical regions around the world (157), and because the epitopes have a different binding profile with different HLA alleles. Thus, population coverage must be taken into a different set of alleles to cover all regions as possible and to get desirable immune response in all individuals within a given population. For that reason, all promising MHC-I and MHC-II epitope candidates were assessed for population coverage against the whole world population. The promising candidates were run against different MHC coverage approaches: class I separate, class II separate, and class I and class II combined, through IEDB population coverage calculation tool at (http://tools.iedb.org/population/) (59, 158).

### 2.6 The Physicochemical properties

The main purpose of vaccination is to induce an immune response after injecting the vaccine into the body. Therefore, it is essential to define the physical and chemical parameters associated with the vaccine. The physicochemical properties of Heat shock 70KDa protein was analyzed using BioEdit sequence alignment editor software Version 7.2.5, and Expasy server (ProtScale and Protparam) (129, 159).

### 2.7 Homology Modelling

The reference sequence of heat shock 70KDa protein was submitted to Raptor X template-based tertiary structure prediction on 29^th^ September 2018 in order to get the 3D structure (160–163). After which, the proposed 3D structure was processed with UCSF chimera 1.13.1 software in order to visualize and allocate the exact sequential location of the selected promiscuous T-cell and B-cell epitope within heat shock 70KDa protein (164–174).

### 2.8 Molecular docking analysis

Molecular docking was performed using Moe 2007(175), the 3D structures of the promiscuous epitopes were predicted by PEP-FOLD (176–179). HLA-C*12:03, HLA-DRB1*01:01 and Immunoglobulin G were chosen as a model for molecular docking to predict the strength of association between the binding site and promiscuous epitopes. The crystal structure of the models were downloaded in a PDB format from the RCSB PDB resource. However, the selected crystal structures were in a complex form with ligands. Thus, to simplify the complex structure all water molecules, hetero groups and ligands were removed by Discovery Studio Visualizer 2.5 (180). Partial charge and energy minimization were applied for ligands and targets. In terms of the identification of the binding groove, the potential binding sites in the crystal structure were recognized using the Alpha Site Finder. Finally, ten independent docking runs were carried out for each Peptide and results were retrieved as binding energies. Best poses for each epitope that displayed lowest binding energies were visualized using UCSF chimera 1.13.1 software (164–174).

## 3. Results

### 3.1 B-cell epitope prediction

With respect to Continuous B-cell epitope prediction yield, In Bepipred Linear Epitope Prediction, the default threshold score of heat shock 70KDa protein to B-cell was given to be 0.249, 298 linear epitopes were predicted. In Emini surface accessibility prediction, the default threshold score of surface accessibility test of the protein was found to be 1.000, 211 epitopes were potentially at the surface by passing the default threshold. In Kolaskar and Tongaonkar antigenicity prediction, the default threshold score of antigenicity was set to be 1.024, fifteen immunogenic epitopes were passed the test. Hence, fifteen linear conserved surface antigenic epitopes were passed all of the above tests; six epitopes out of all were thought to be the promising B-cell epitopes that are able to evoke B lymphocyte for their efficient physiochemical properties and length (**ANYVQASEK, NYVQASEK, EKPKNVNPVI, EVEKEEEVTV, KSVEKPAS,** and **TPQQPPAQ)**. With regard to discontinuous B-cell epitope prediction yield, seven promising discontinuous epitopes were defined from the modeled 3D after submission to ElliPro prediction tool; epitopes were predicted to be located on the surface of the protein indicating quick recognition by host immune system.

**Table 1.** List of the fifteen linear conserved surface antigenic epitopes with their surface accessibility score and antigenicity score.

| Start | End | Linear epitopes <sup>a</sup> | Length | Surface score <sup>b</sup> | Antigenicity score <sup>c</sup> |
| --- | --- | --- | --- | --- | --- |
| 79 | 84 | PEVEEY | 6 | 1.748 | 1.027 |
| 592 | 600 | ANYVQASEK* | 9 | 1.243 | 1.028 |
| 593 | 600 | NYVQASEK* | 8 | 1.625 | 1.024 |
| 594 | 600 | YVQASEK | 7 | 1.335 | 1.059 |
| 595 | 600 | VQASEK | 6 | 1.128 | 1.042 |
| 468 | 474 | VKSVEKP | 7 | 1.118 | 1.079 |
| 705 | 716 | QSEKPKNVNPVI | 12 | 1.3 | 1.028 |
| 706 | 716 | SEKPKNVNPVI | 11 | 1.002 | 1.029 |
| 707 | 716 | EKPKNVNPVI* | 10 | 1.003 | 1.031 |
| 504 | 513 | EVEKEEEVTV* | 10 | 1.115 | 1.024 |
| 735 | 741 | ILNKPKP | 7 | 1.257 | 1.024 |
| 735 | 745 | ILNKPKPKPKV | 11 | 1.844 | 1.043 |
| 735 | 746 | ILNKPKPKPKVT | 12 | 1.995 | 1.032 |
| 469 | 476 | KSVEKPAS* | 8 | 1.544 | 1.031 |
| 749 | 756 | TPQQPPAQ* | 8 | 2.997 | 1.026 |

\*Top promising epitopes for their ideal length and physiochemical properties
<sup>a</sup> Bepipred Linear default threshold value 0.249
<sup>b</sup> Emini surface accessibility default threshold value 1.000
<sup>c</sup> Kolaskar and Tongaonkar antigenicity default threshold value 1.024

**Table 2.**
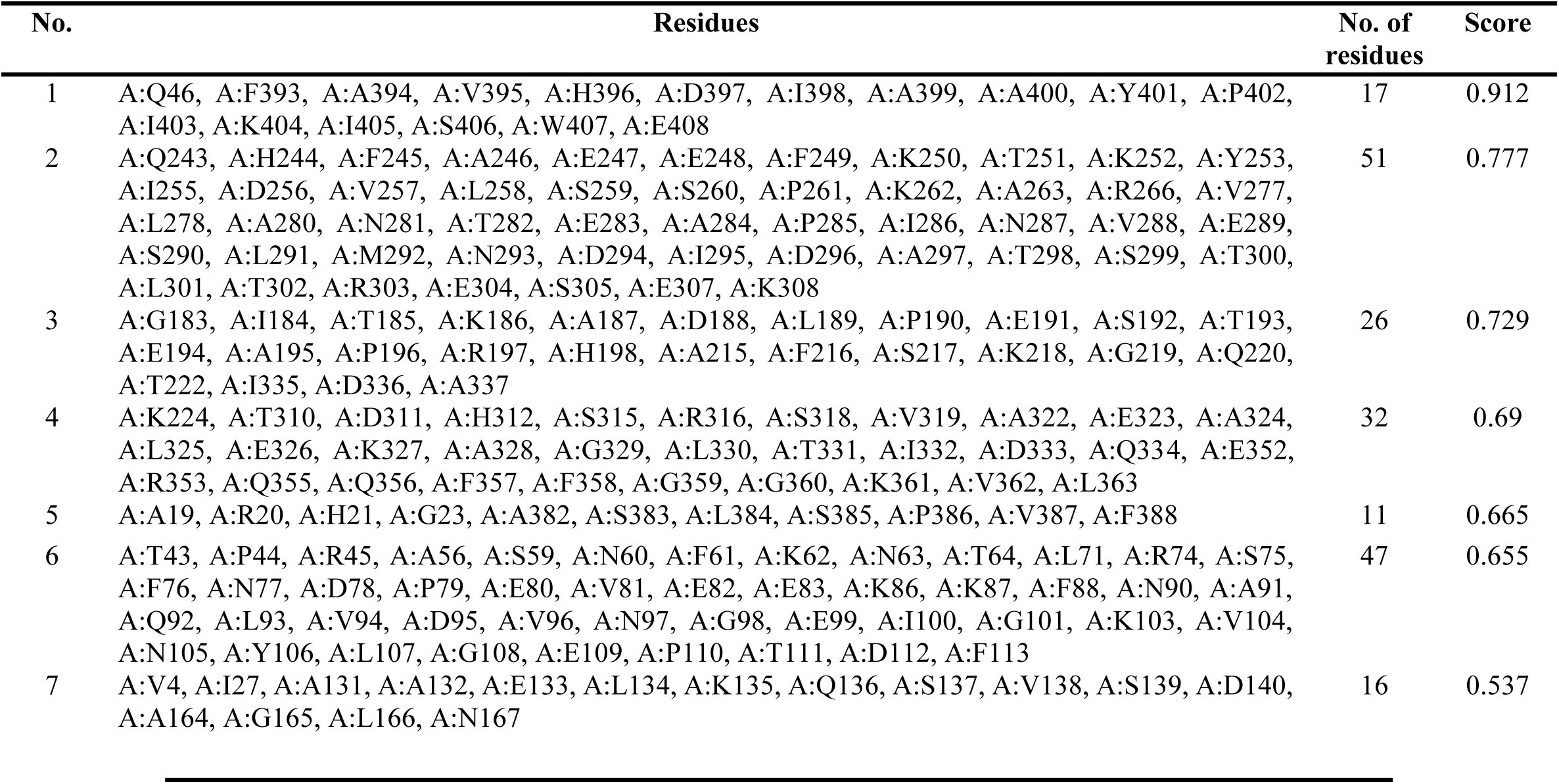
List of Predicted discontinuous B-cell epitopes by ElliPro prediction tool with number of Residues and their scores.

| No. | Residues | No. of residues | Score |
| --- | --- | --- | --- |
| 1 | A:Q46, A:F393, A:A394, A:V395, A:H396, A:D397, A:I398, A:A399, A:A400, A:Y401, A:P402, A:I403, A:K404, A:I405, A:S406, A:W407, A:E408 | 17 | 0.912 |
| 2 | A:Q243, A:H244, A:F245, A:A246, A:E247, A:E248, A:F249, A:K250, A:T251, A:K252, A:Y253, A:I255, A:D256, A:V257, A:L258, A:S259, A:S260, A:P261, A:K262, A:A263, A:R266, A:V277, A:L278, A:A280, A:N281, A:T282, A:E283, A:A284, A:P285, A:I286, A:N287, A:V288, A:E289, A:S290, A:L291, A:M292, A:N293, A:D294, A:I295, A:D296, A:A297, A:T298, A:S299, A:T300, A:L301, A:T302, A:R303, A:E304, A:S305, A:E307, A:K308 | 51 | 0.777 |
| 3 | A:G183, A:I184, A:T185, A:K186, A:A187, A:D188, A:L189, A:P190, A:E191, A:S192, A:T193, A:E194, A:A195, A:P196, A:R197, A:H198, A:A215, A:F216, A:S217, A:K218, A:G219, A:Q220, A:T222, A:I335, A:D336, A:A337 | 26 | 0.729 |
| 4 | A:K224, A:T310, A:D311, A:H312, A:S315, A:R316, A:S318, A:V319, A:A322, A:E323, A:A324, A:L325, A:E326, A:K327, A:A328, A:G329, A:L330, A:T331, A:I332, A:D333, A:Q334, A:E352, A:R353, A:Q355, A:Q356, A:F357, A:F358, A:G359, A:G360, A:K361, A:V362, A:L363 | 32 | 0.69 |
| 5 | A:A19, A:R20, A:H21, A:G23, A:A382, A:S383, A:L384, A:S385, A:P386, A:V387, A:F388 | 11 | 0.665 |
| 6 | A:T43, A:P44, A:R45, A:A56, A:S59, A:N60, A:F61, A:K62, A:N63, A:T64, A:L71, A:R74, A:S75, A:F76, A:N77, A:D78, A:P79, A:E80, A:V81, A:E82, A:E83, A:K86, A:K87, A:F88, A:N90, A:A91, A:Q92, A:L93, A:V94, A:D95, A:V96, A:N97, A:G98, A:E99, A:I100, A:G101, A:K103, A:V104, A:N105, A:Y106, A:L107, A:G108, A:E109, A:P110, A:T111, A:D112, A:F113 | 47 | 0.655 |
| 7 | A:V4, A:I27, A:A131, A:A132, A:E133, A:L134, A:K135, A:Q136, A:S137, A:V138, A:S139, A:D140, A:A164, A:G165, A:L166, A:N167 | 16 | 0.537 |

**Figure 2.**
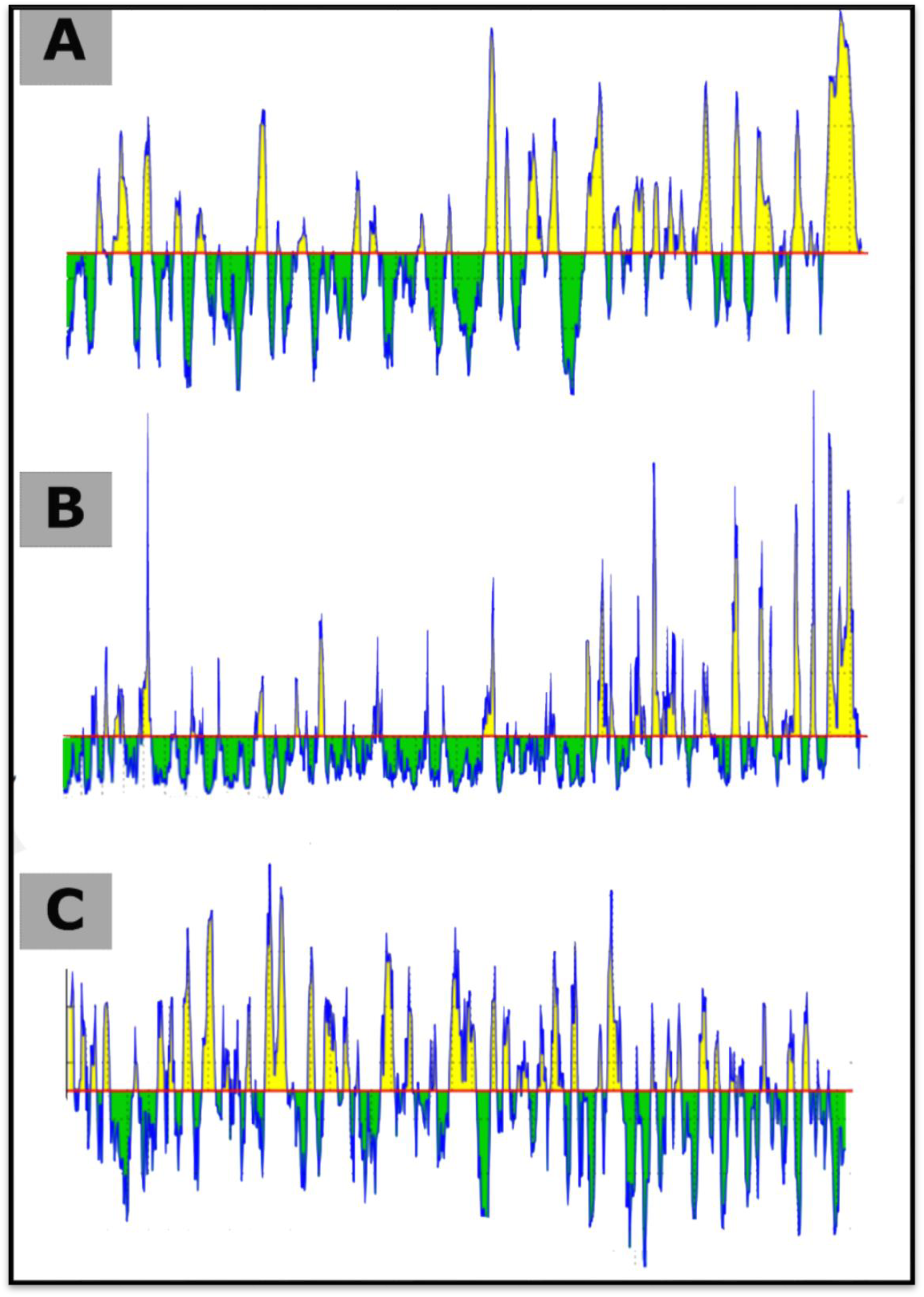
Illustrates the spectrums of the linear conserved surface immunogenic B-cell epitopes. **(A)** Bepipred Linear Epitope Prediction, the yellow spectrums above and at a cut-off of 0.249 (red line) represent the linear epitopes while the green spectrums exemplify the non-linear epitopes. **(B)** Emini surface accessibility prediction, the yellow spectrums above and at a cut-off of 1.000 (red line) illustrate epitopes on the surface whereas green spectrums represent epitopes that aren’t on the surface. **(C)** Kolaskar and Tongaonkar antigenicity prediction, the yellow spectrums above and at a cut-off of 1.024 (red line) represent the immunogenic epitopes while green spectrums demonstrate the non-immunogenic or zero fold epitopes.

**Figure 3.**
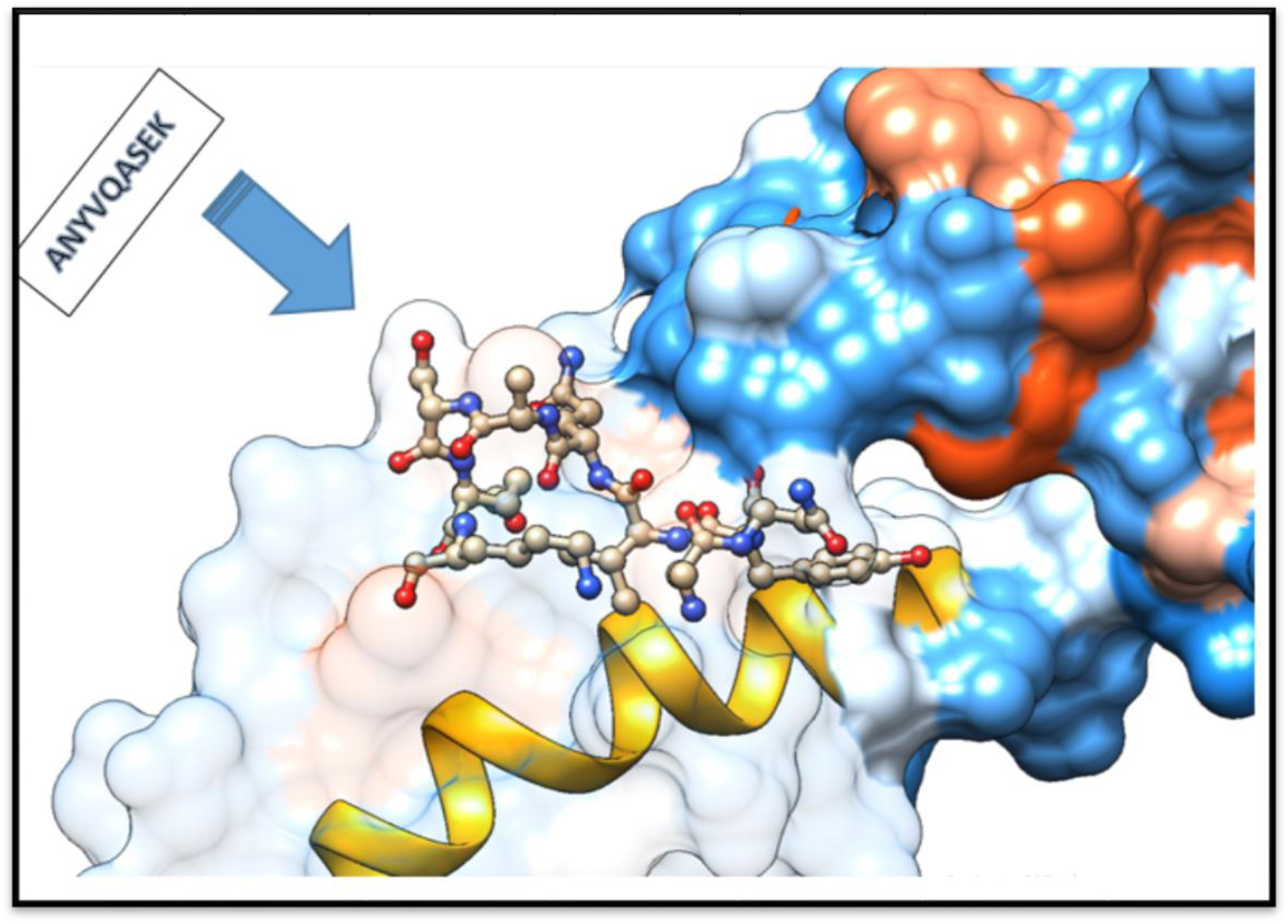
Sequential position of **ANYVQASEK** as a promising B-cell epitope within the 3D structure of heat shock 70KDa protein of *C. neoformans* using UCSF chimera 1.13.1 software.

**Figure 4.**
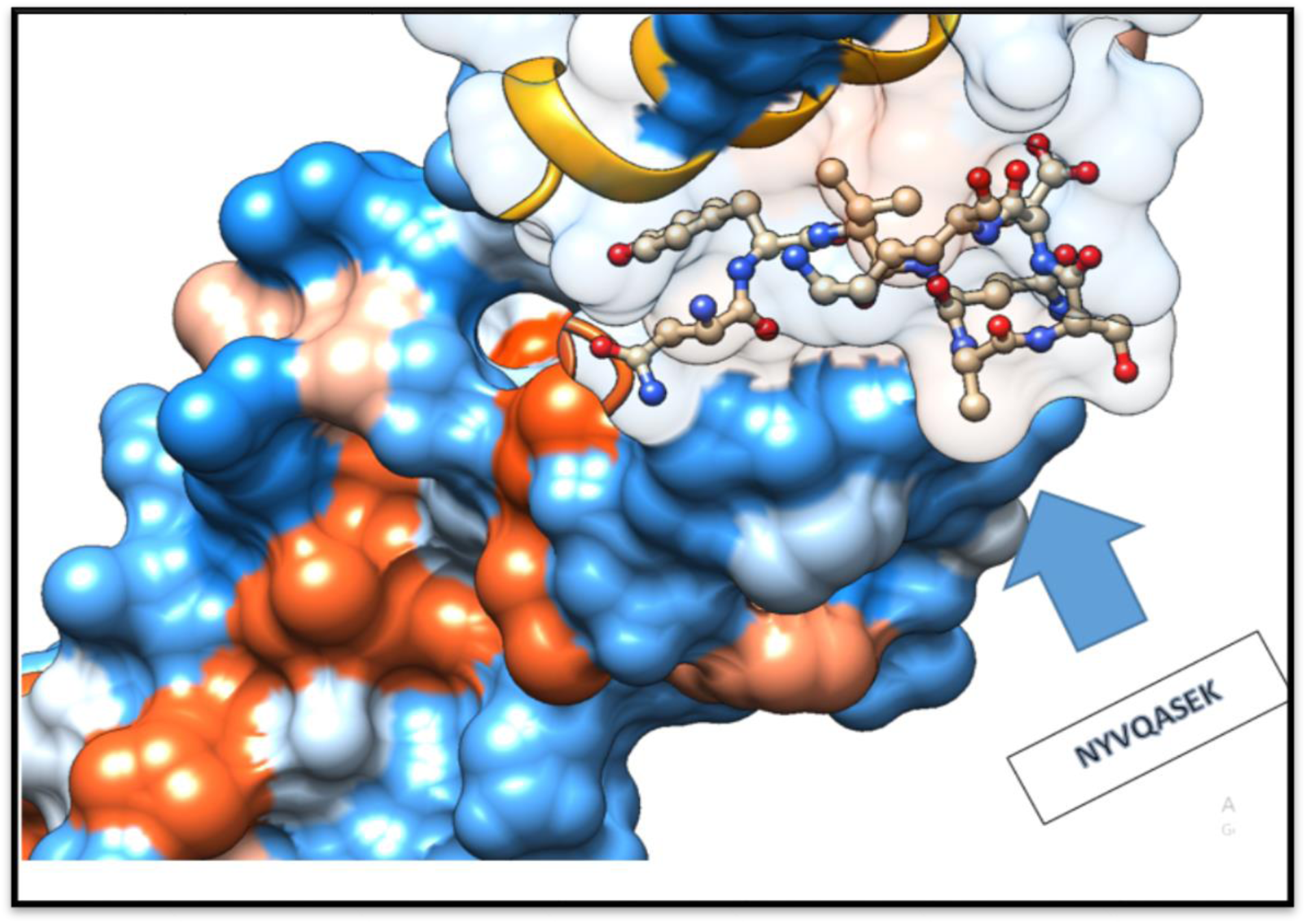
Sequential position of **NYVQASEK** as a promising B-cell epitope within the 3D structure of heat shock 70KDa protein of *C. neoformans* using UCSF chimera 1.13.1 software.

**Figure 5.**
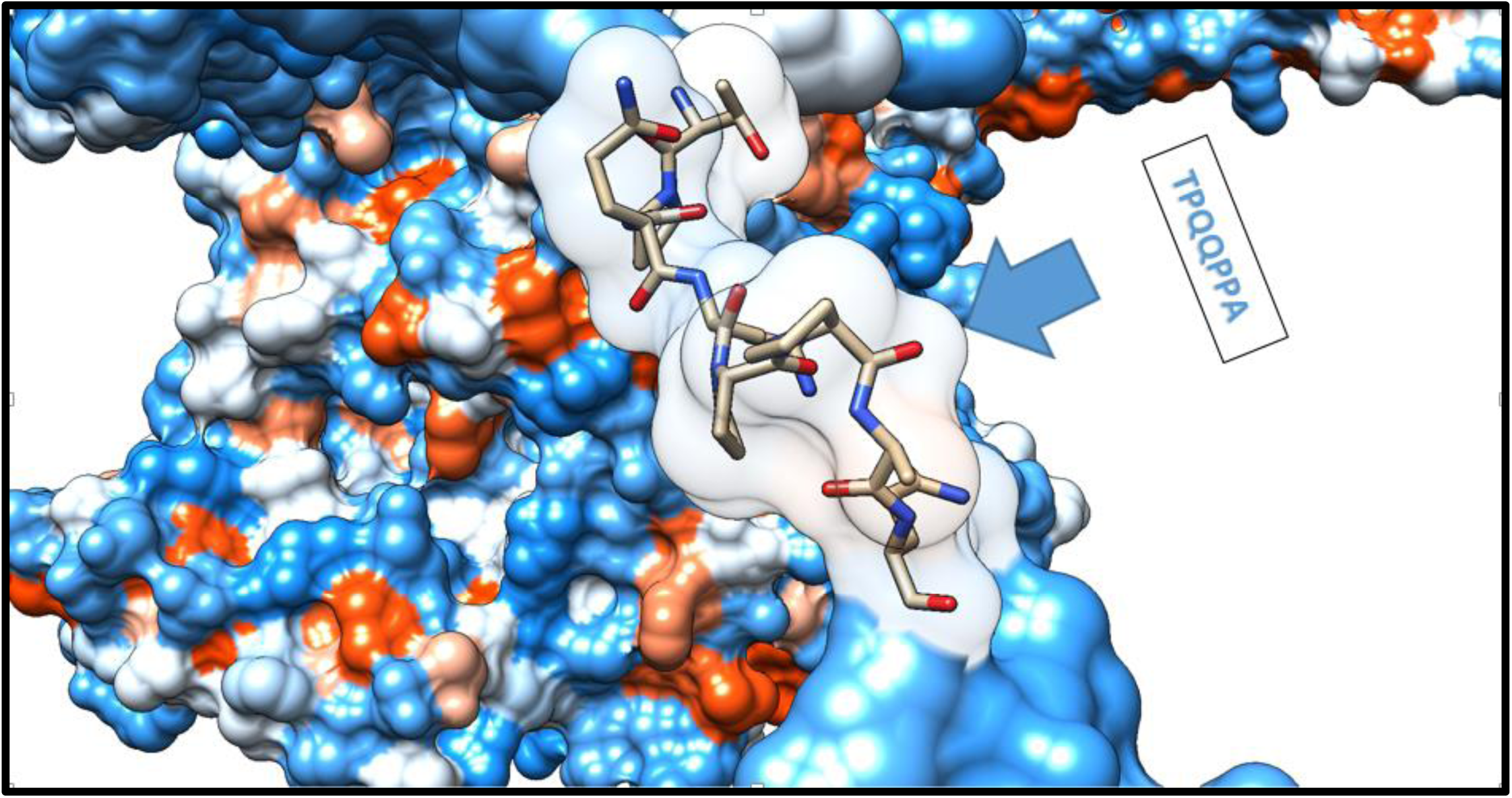
Sequential position of **TPQQPPAQ** as a promising B-cell epitope within the 3D structure of heat shock 70KDa protein of *C. neoformans* using UCSF chimera 1.13.1 software.

**Figure 6.**
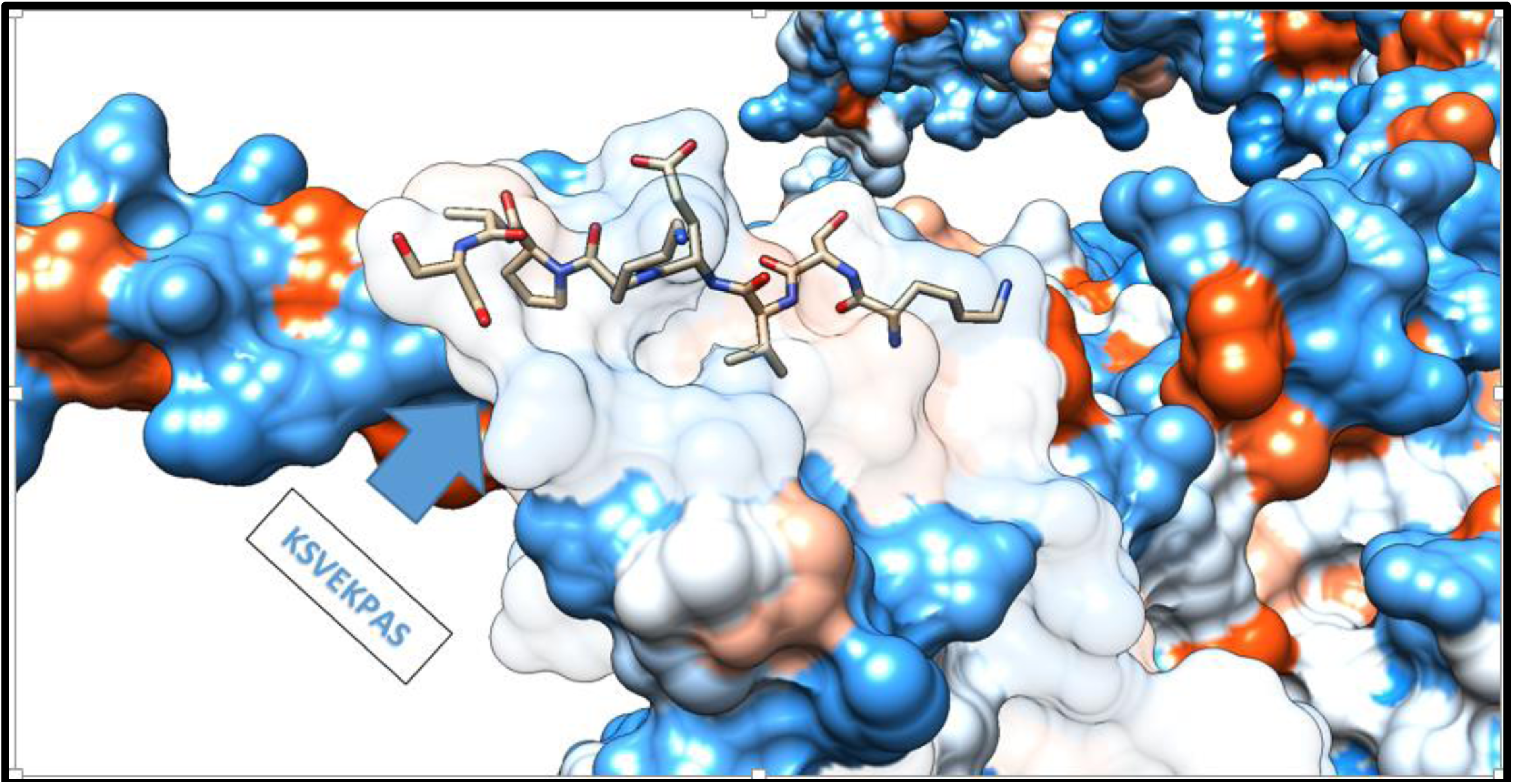
Sequential position of **KSVEKPAS** as a promising B-cell epitope within the 3D structure of heat shock 70KDa protein of *C. neoformans* using UCSF chimera 1.13.1 software.

**Figure 7.**
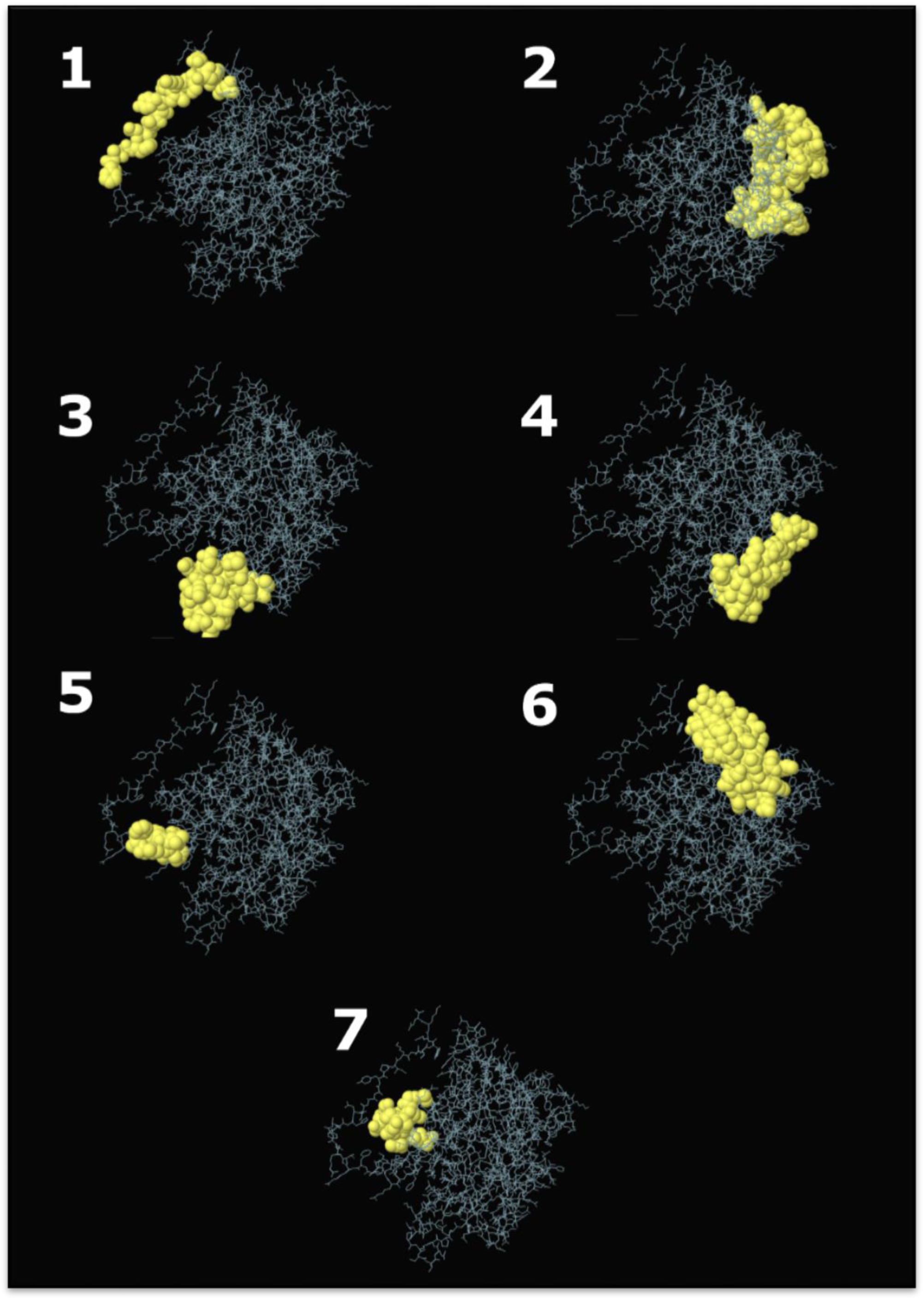
Three-dimensional representation of discontinuous epitopes (1–7) of the highest immunogenic heat shock 70KDa protein of *C. neoformans* using ElliPro prediction tool. The epitopes are depicted in yellow surface, and the bulk of the heat shock 70KDa protein is depicted in grey sticks.

### 3.2 Prediction of MHC-I binding profile for conserved epitopes

213 epitopes were anticipated to interact with different MHC-1 alleles. The core epitopes **YVYDTRGKL** was noticed to be the dominant binder with 9 alleles (HLA-A*02:06, HLA-A*68:02, HLA-B*07:02, HLA-C*03:03, HLA-C*06:02, HLA-C*07:01, HLA-C*12:03, HLA-C*14:02, HLA-C*15:02), followed by **LTFYRQGAF, RATPSLVSF** and **FTQLVAAYL** are believed to bind with 7 alleles for each.

**Table 2.**
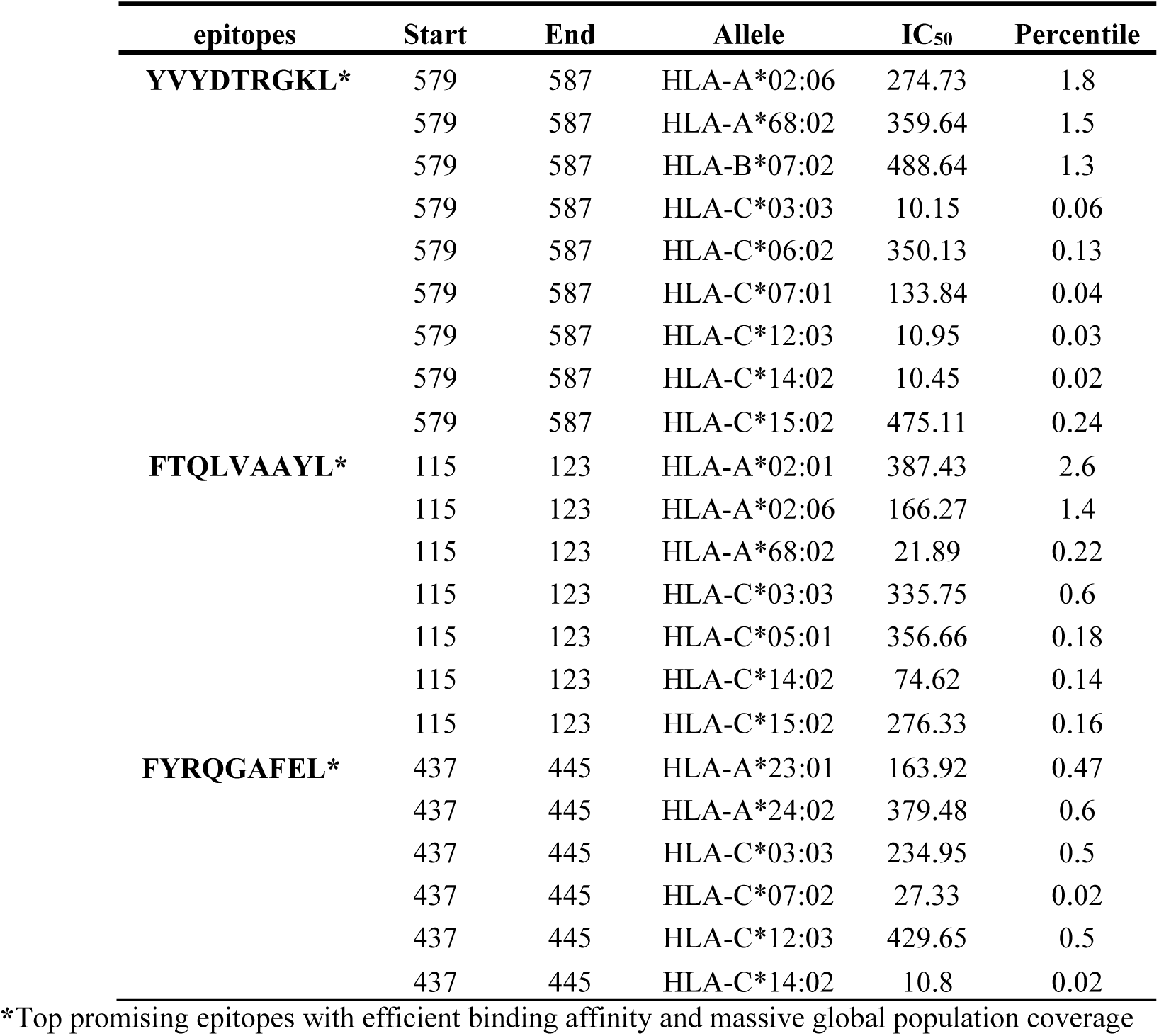
List of promising epitopes that had a good binding affinity with MHC-I alleles in terms of IC**_50_** and Percentile rank.

**Figure 8.**
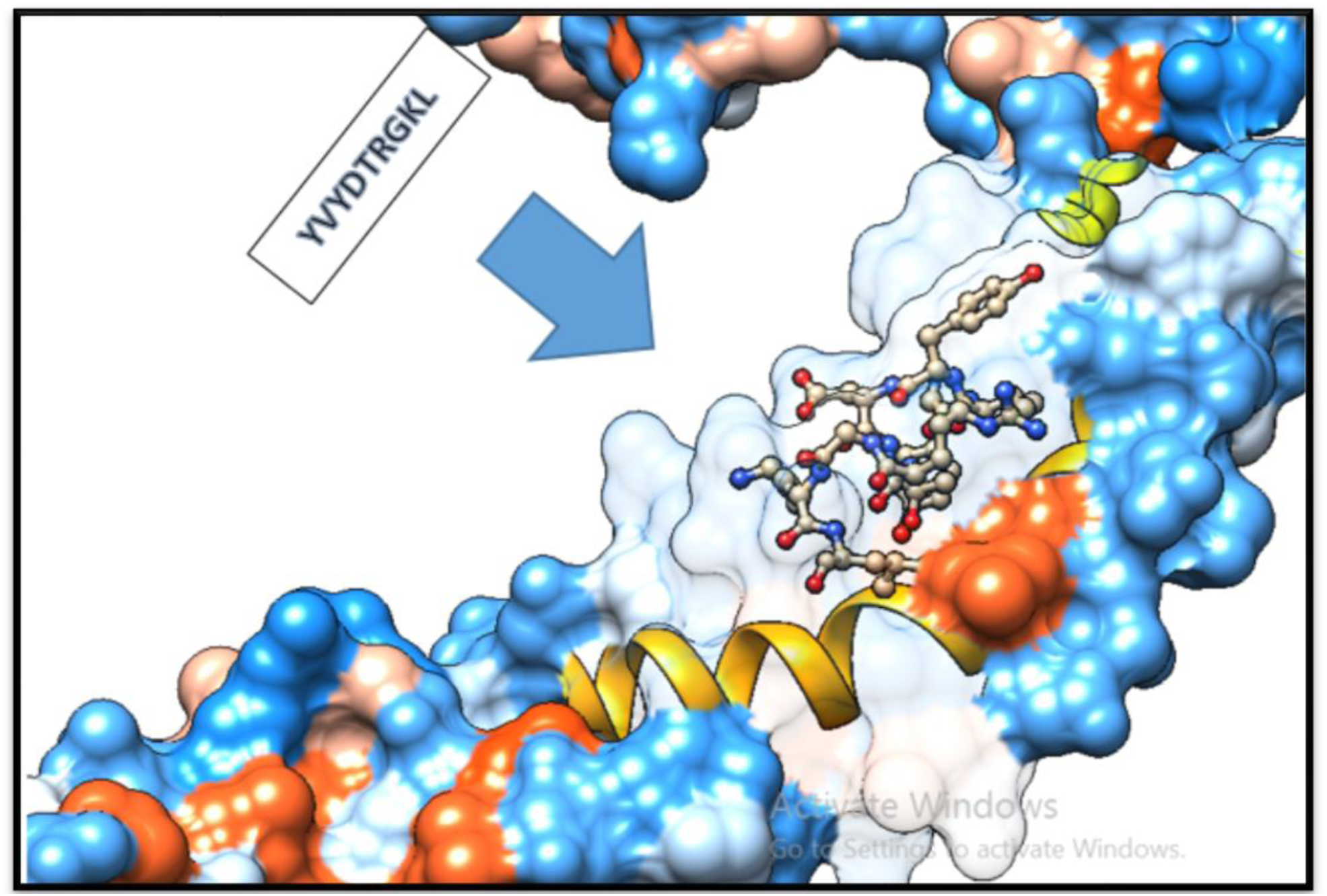
Structural location of **YVYDTRGKL** as a promising MHC-I binder, with massive population coverage, within the 3D structure of *C. neoformans’s* heat shock 70KDa protein using UCSF chimera 1.13.1 software.

**Figure 9.**
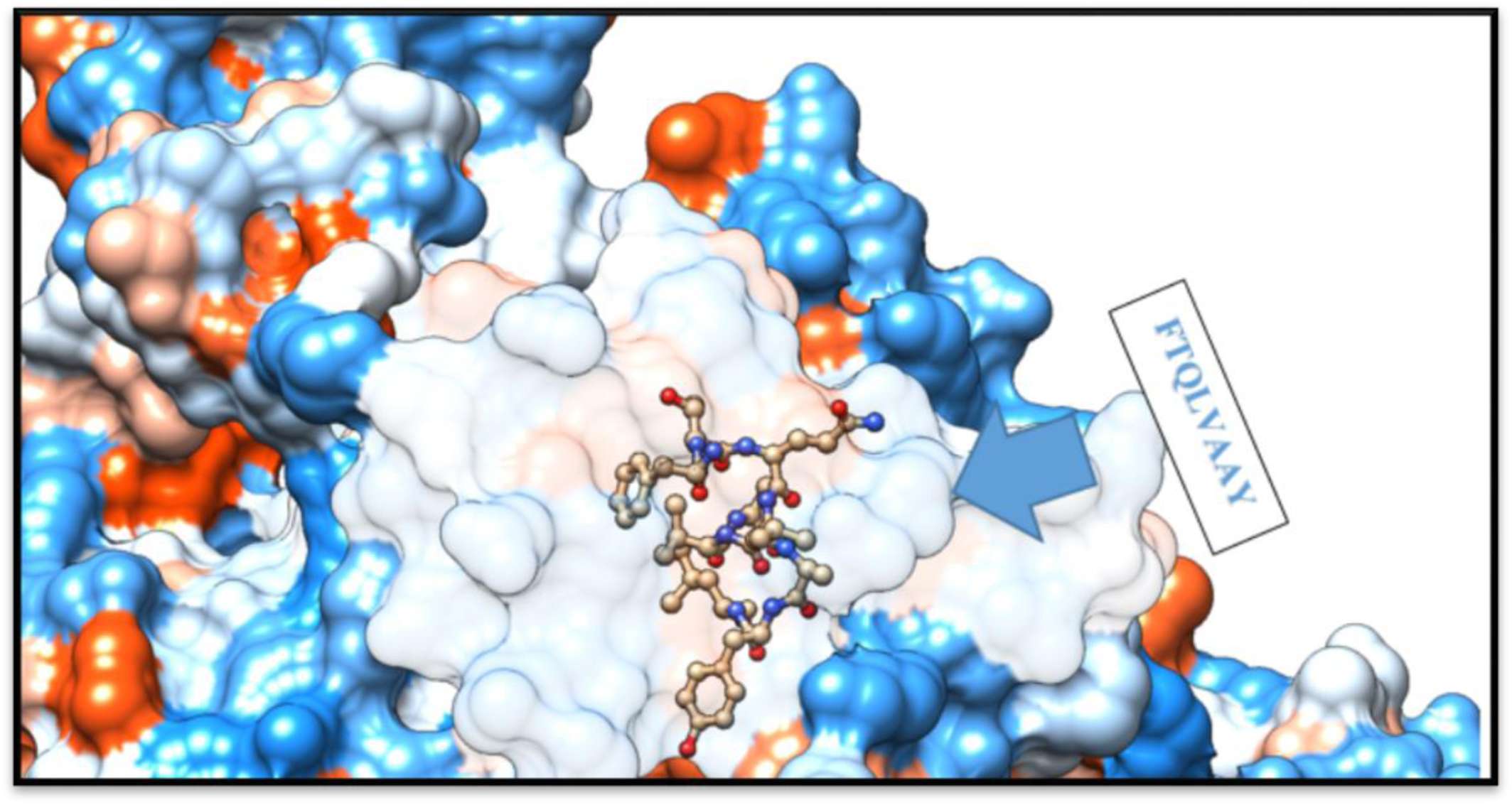
Structural location of **FTQLVAAYL** as a promising MHC-I binder, with massive population coverage, within the 3D structure of *C. neoformans’s* heat shock 70KDa protein using UCSF chimera 1.13.1 software.

**Figure 10.**
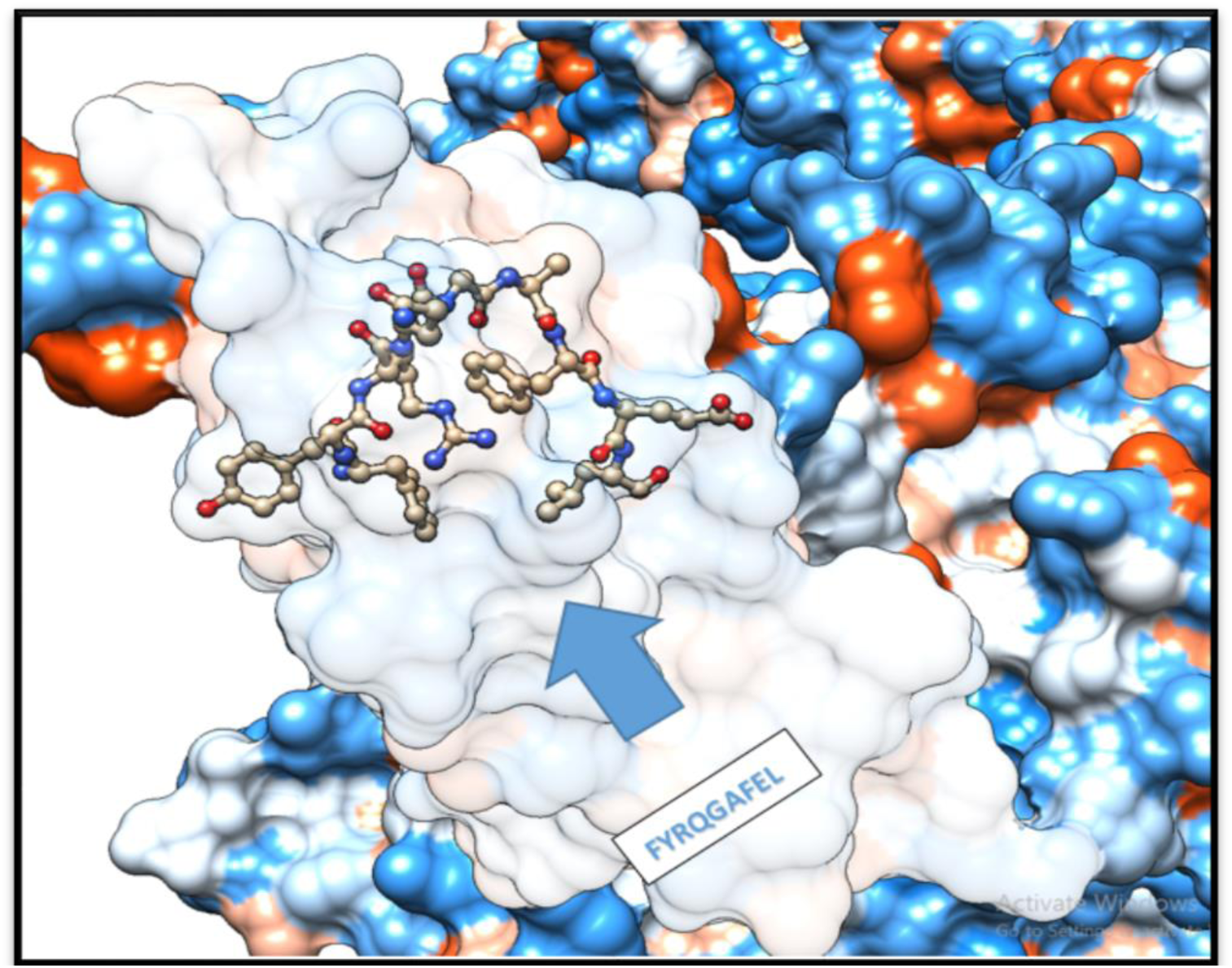
Structural location of **FYRQGAFEL** as a promising MHC-II epitope, with massive population coverage, within the 3D structure of *C. neoformans’s* heat shock 70KDa protein using UCSF chimera 1.13.1 software.

### 3.3 Prediction of MHC-II binding profile for conserved epitopes

156 conserved predicted epitopes found to interact with MHC-II alleles. The core epitopes **FDYALVQHF** is thought to be the top binder as it interacts with eleven alleles; (HLA-DPA1*01:03, HLA-DPB1*02:01, HLA-DPA1*02:01, HLA-DPB1*01:01, HLA-DRB1*01:01, HLA-DRB1*03:01, HLA-DRB1*04:05, HLA-DRB1*07:01, HLA-DRB1*09:01, HLA-DRB1*11:01, HLA-DRB5*01:01). Followed by **FYRQGAFEL** and **FFGGKVLNF** are believed to bind with 10 alleles for each.

**Table 3.**
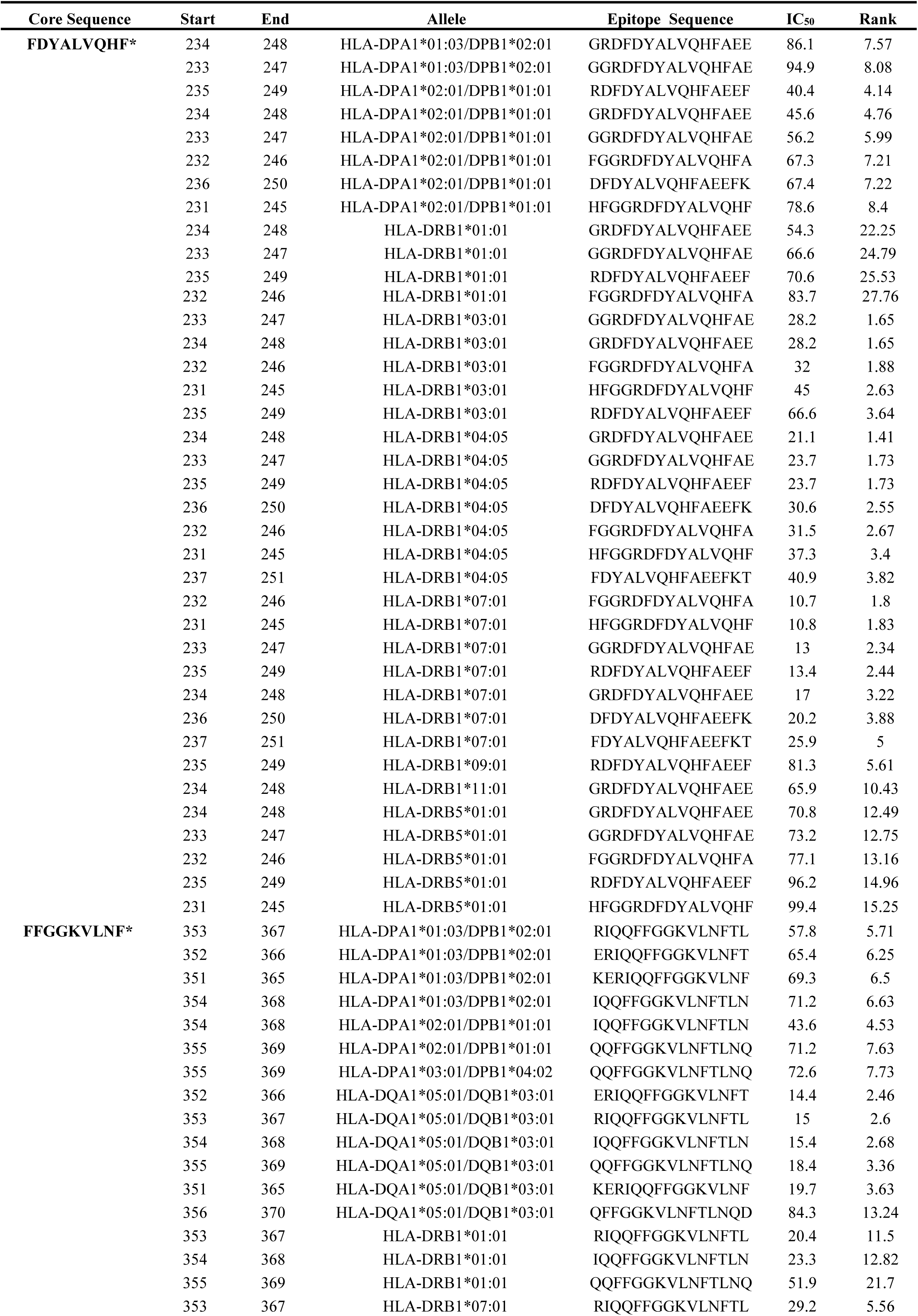
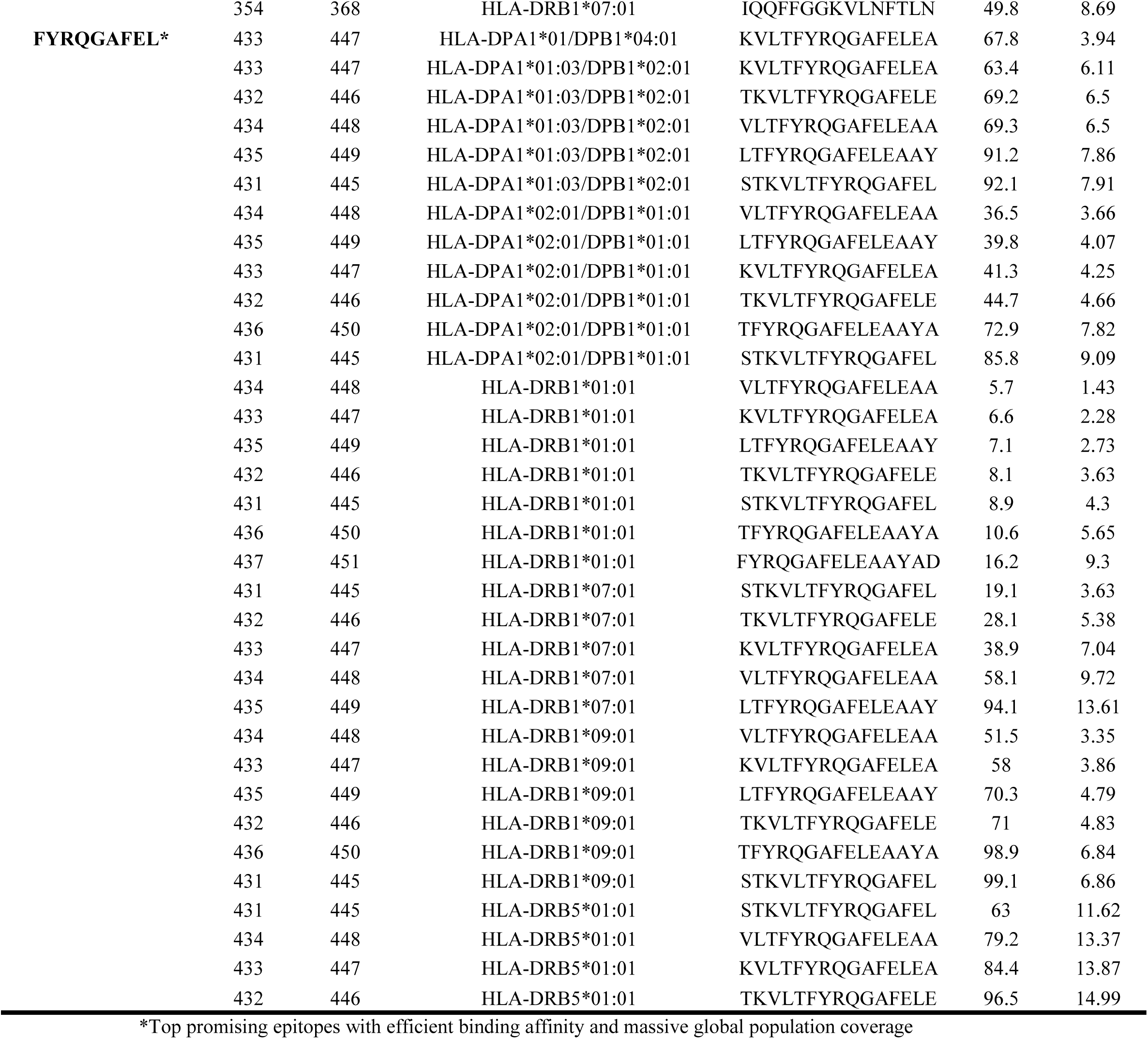
List of the three promising epitopes core sequence that had a strong binding affinity with MHC-II in terms of IC50 and Percentile Ranks.

**Figure 11.**
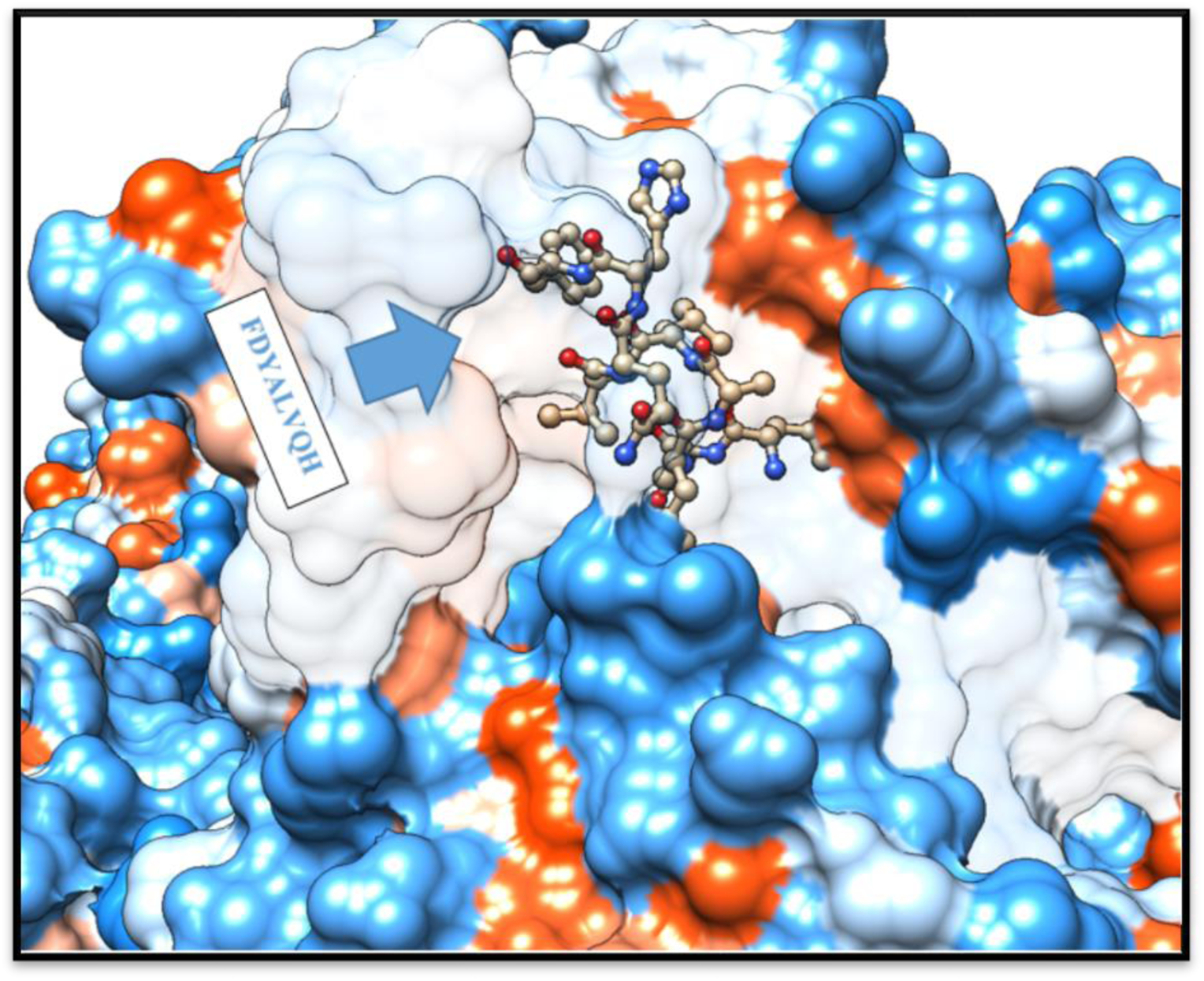
Structural location of **FDYALVQHF** as a promising MHC-II epitope, with massive population coverage, within the 3D structure of *C. neoformans’s* heat shock 70KDa protein using UCSF chimera 1.13.1 software.

**Figure 12.**
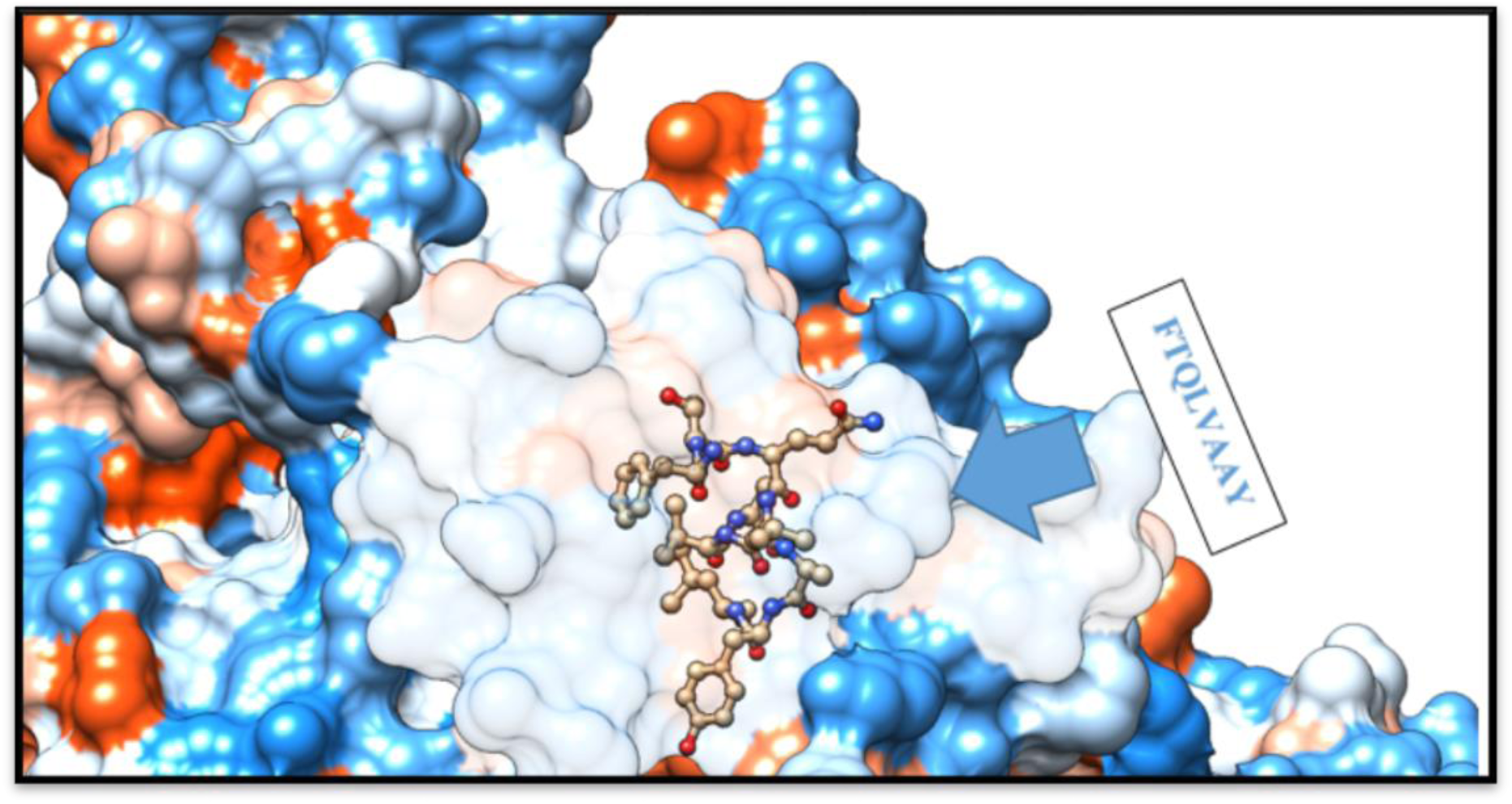
Structural location of **FTQLVAAYL** as a promising MHC-II epitope within the 3D structure of *C. neoformans’s* heat shock 70KDa protein using UCSF chimera 1.13.1 software.

**Figure 13.**
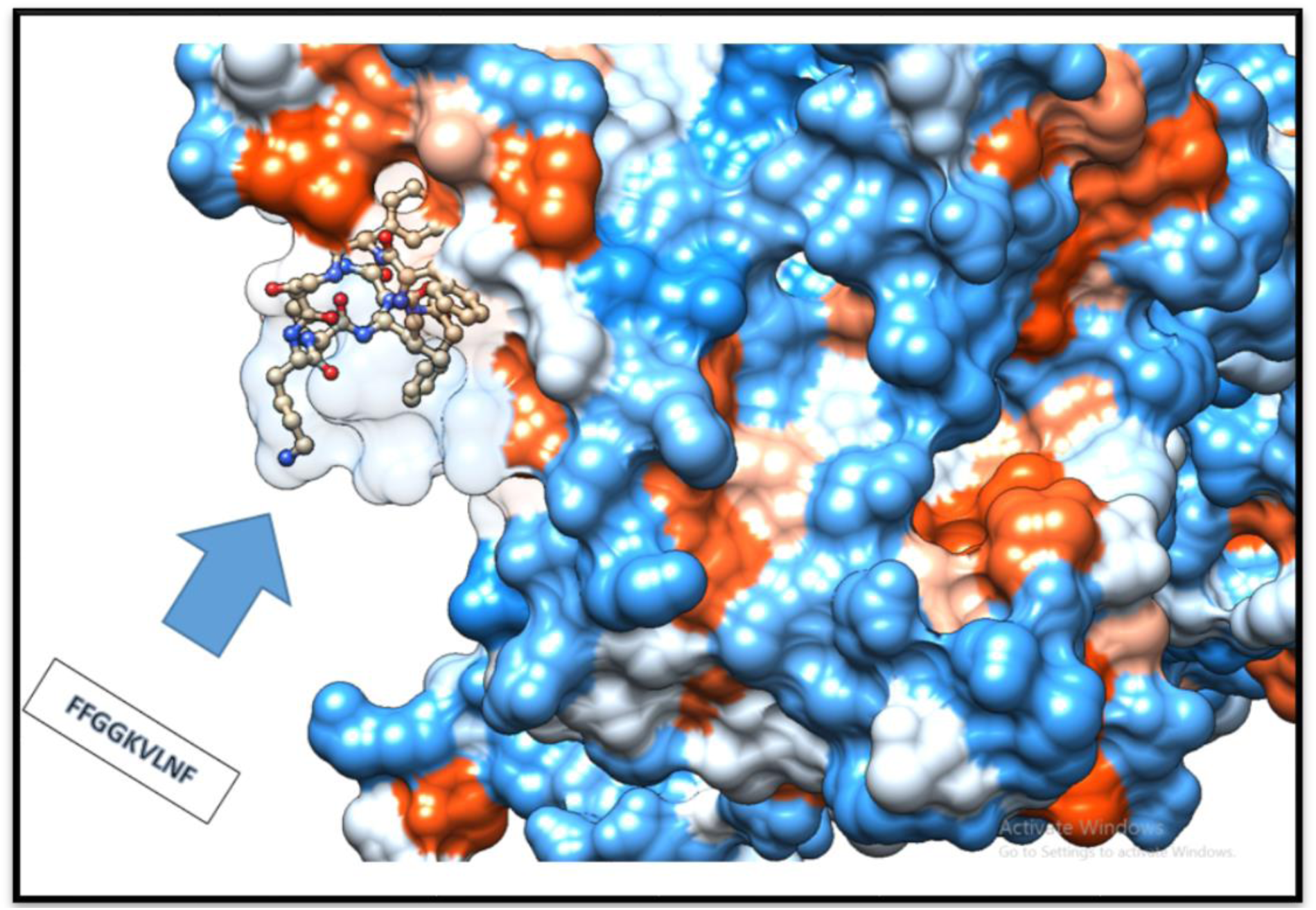
Structural location of **FFGGKVLNF** as a promising MHC-II epitope, with massive population coverage, within the 3D structure of *C. neoformans’s* heat shock 70KDa protein using UCSF chimera 1.13.1 software.

### 3.5 Physiochemical parameters

The protein length was found to be 773 amino acids. MW, and pI parameters were calculated as 85.69KDa and 5.12, respectively. The pI value indicates that the protein is acidic in nature. The total numbers of negatively and positively charged residues were 124 and 98, correspondingly. Extinction coefficient of protein at 280 nm was measured 67520M^−1^ cm^−1^ in water. The half-life of vaccine was predicted to be 30 hours in (mammalian reticulocytes, in vitro),>20 hours in (yeast, in vivo) and >10 hours in (*Escherichia coli*, in vivo). The Instability index was computed to be 35.99, which indicates the thermostability. The aliphatic index and the GRAVY value of vaccine were determined 85.80 and (−0.416), respectively. The high aliphatic index shows that the protein is stable in a wide range of temperatures, and the negative GRAVY value shows protein hydrophilicity and a better interaction with the surrounding water molecules.

**Table 4.**
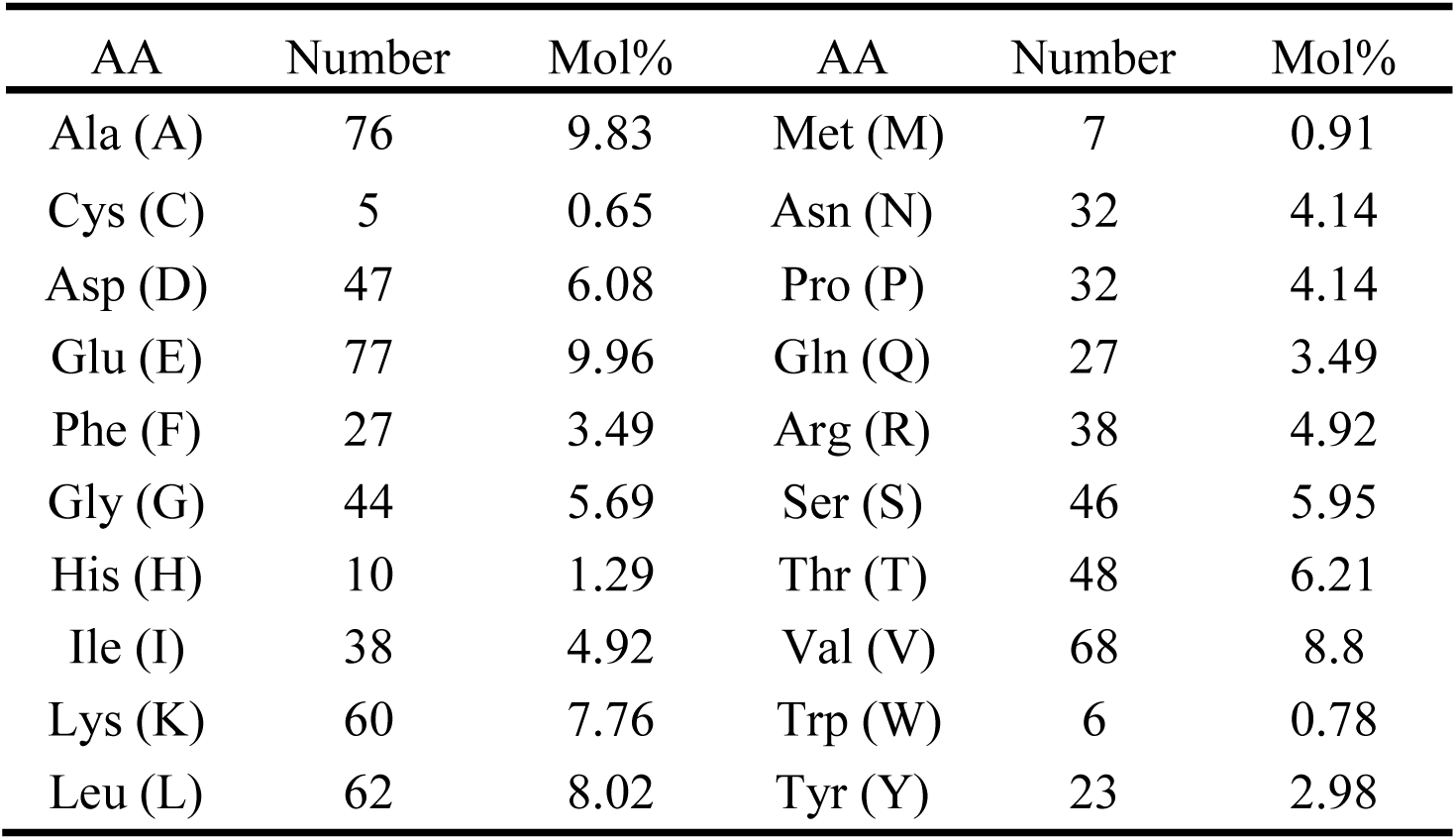
List of amino acid that formed heat shock 70KDa protein, their sequential location, and molar percentage (Mol %)

| AA | Number | Mol% | AA | Number | Mol% |
| --- | --- | --- | --- | --- | --- |
| Ala (A) | 76 | 9.83 | Met (M) | 7 | 0.91 |
| Cys (C) | 5 | 0.65 | Asn (N) | 32 | 4.14 |
| Asp (D) | 47 | 6.08 | Pro (P) | 32 | 4.14 |
| Glu (E) | 77 | 9.96 | Gln (Q) | 27 | 3.49 |
| Phe (F) | 27 | 3.49 | Arg (R) | 38 | 4.92 |
| Gly (G) | 44 | 5.69 | Ser (S) | 46 | 5.95 |
| His (H) | 10 | 1.29 | Thr (T) | 48 | 6.21 |
| Ile (I) | 38 | 4.92 | Val (V) | 68 | 8.8 |
| Lys (K) | 60 | 7.76 | Trp (W) | 6 | 0.78 |
| Leu (L) | 62 | 8.02 | Tyr (Y) | 23 | 2.98 |

**Figure 14.**
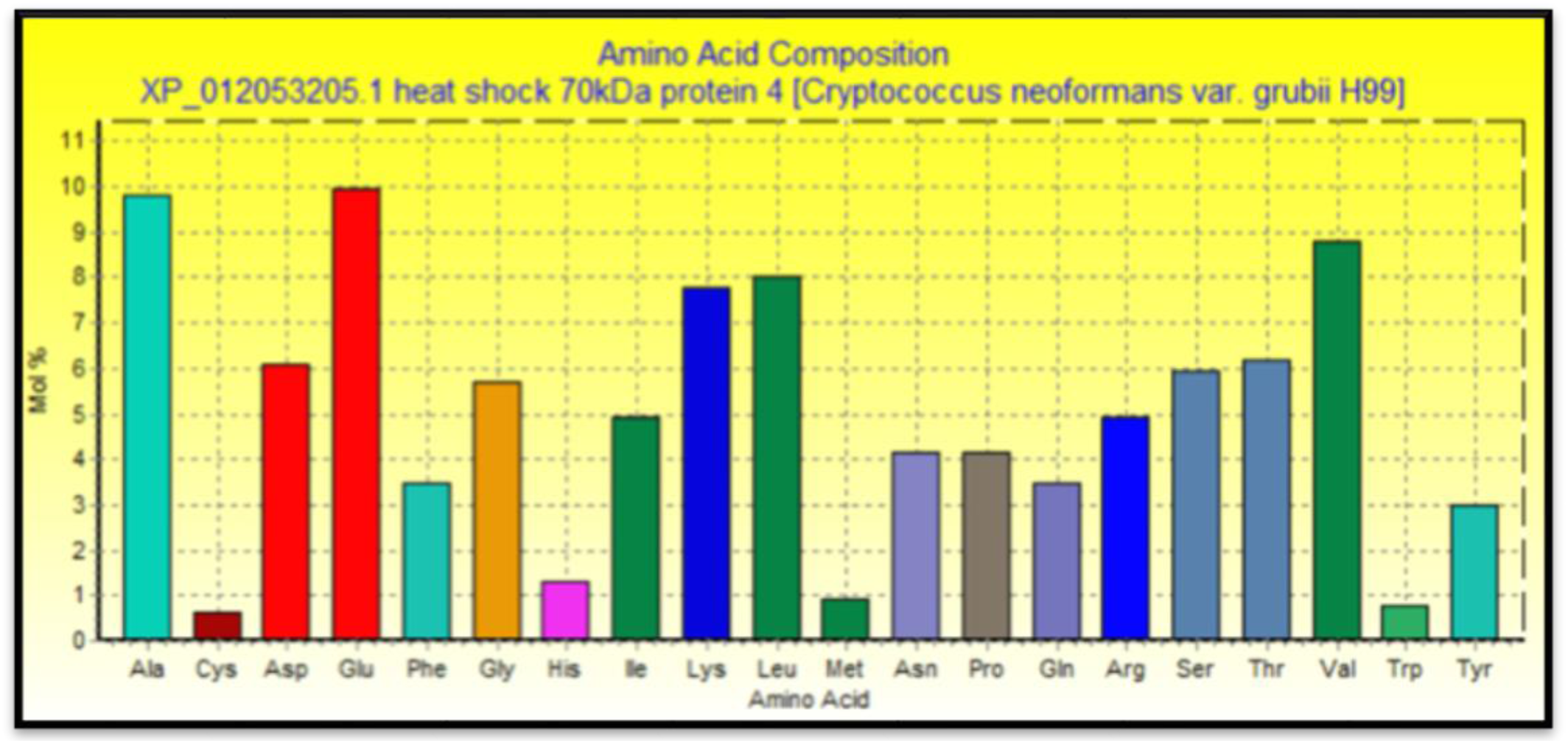
The graph shows the Amino Acid Composition (Mol %) of heat shock 70KDa protein using BioEdit sequence alignment editor software Version 7.2.5.

**Figure 15.**
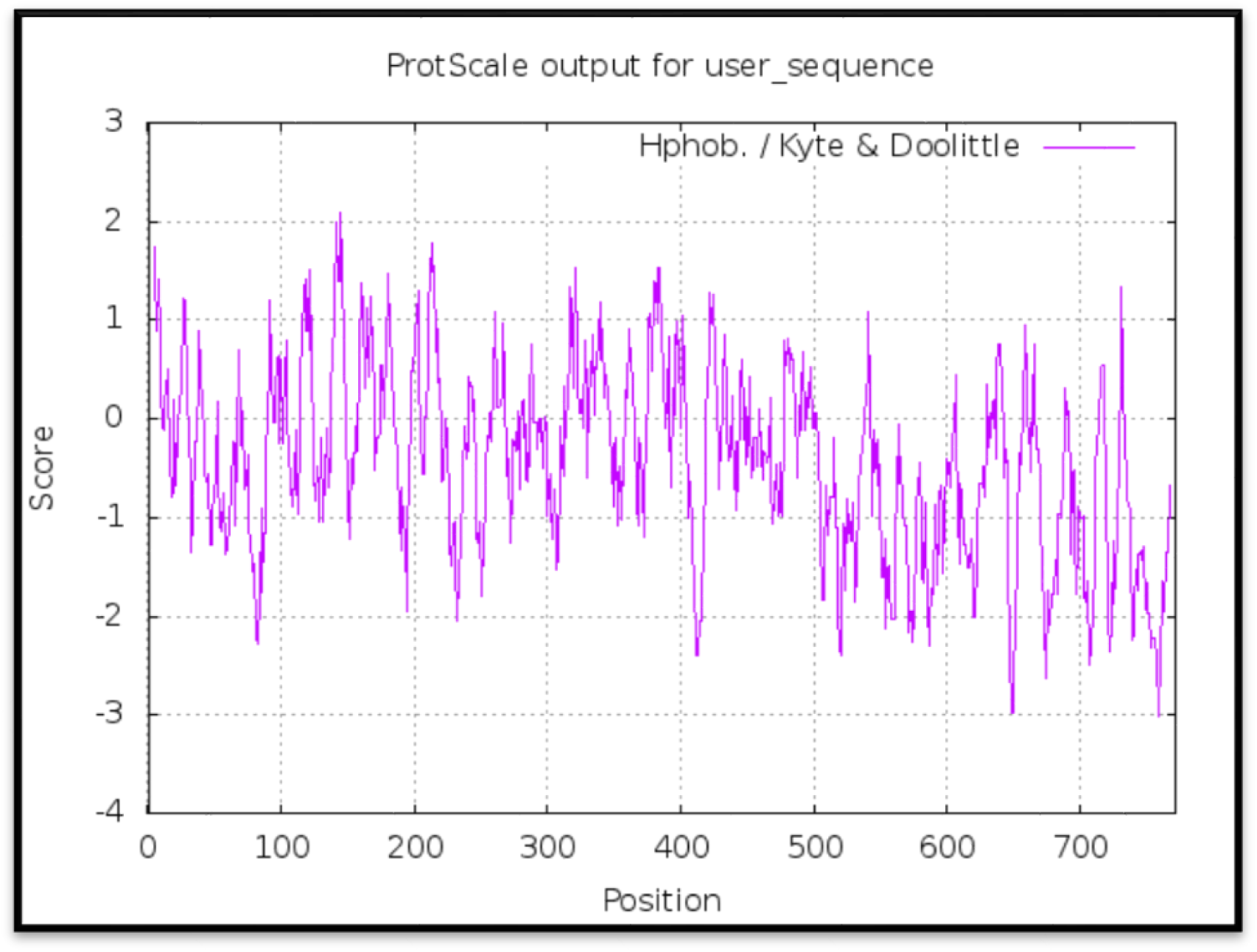
The graph shows the Hydrophobicity scale of heat shock 70KDa protein using ProtScale server.

### 3.4 Population coverage analysis

Population coverage test was performed to compute the world coverage of epitopes that bind to separate MHC-I alleles, MHC-II alleles for each, and combined MHC-I and MHC-II. In addition to, to sort out the most predominant promising epitopes for each coverage mode through IEDB population coverage analysis tool.

#### 3.4.1 Population coverage for isolated MHC-I & MHC-II

When it comes to MHC class I, three epitopes are given to interact with most frequent MHC class I alleles (**YVYDTRGKL, FYRQGAFEL,** and **FTQLVAAYL)**; represent a considerable coverage against the whole world residents. The maximum population coverage percentage over these epitopes in the World was found to be 60.93% for **YVYDTRGKL.**

In the case MHC class II, three epitopes were assumed to interact with most frequent MHC class II alleles (**FFGGKVLNF, FYRQGAFEL,** and **FDYALVQHF**); infer a massive global coverage. The most abundant population coverage percentage of these epitopes in the World was awarded to **FFGGKVLNF** with percentage of 98.02%.

What is interesting in this test is that the population coverage analysis result for the most abundant binders to MHC-I and MHC-II alleles for each in combined form; exhibit an exceptional coverage with the percent of 90.64%, 99.3%, respectively.

**Table 5.** Global Population coverage of promising epitopes in isolated MHC class I and II

| Core epitope | Coverage<br>World<br>class I | Total<br>HLA<br>hits | Core epitopes | Coverage<br>World<br>Class II | Total<br>HLA<br>hits |
| --- | --- | --- | --- | --- | --- |
| **YVYDTRGKL\*** | 60.93% | 9 | **FFGGKVLNF\*** | 98.02% | 10 |
| **FYRQGAFEL\*** | 55.50% | 6 | **FYRQGAFEL\*** | 95.39% | 10 |
| **FTQLVAAYL\*** | 55.41% | 7 | **FDYALVQHF\*** | 95.38% | 11 |
| FACASLSPV | 55.16% | 6 | VVFGTANPI | 77.79% | 6 |
| LLSRVSVPL | 53.01% | 5 | FTQLVAAYL | 61.30% | 8 |
| YADPASLPK | 45.71% | 6 | FKNTVGSLK | 36.46% | 5 |
| RATPSLVSF | 38.17% | 7 | IAGLNALRL | 56.08% | 7 |
| GIMNFEGAY | 36.03% | 6 | LKRLIGRSF | 42.52% | 5 |
| LTFYRQGAF | 35.53% | 7 | IVKVKARLN | 39.32% | 5 |
| FVDVGHSDY | 34.93% | 5 | LREALNTYL | 36.70% | 5 |
| **Peptide set** | 47.04% |  | **Peptide set** | 63.90% |  |

\*Top promising epitopes with massive population coverage

**Figure 16.**
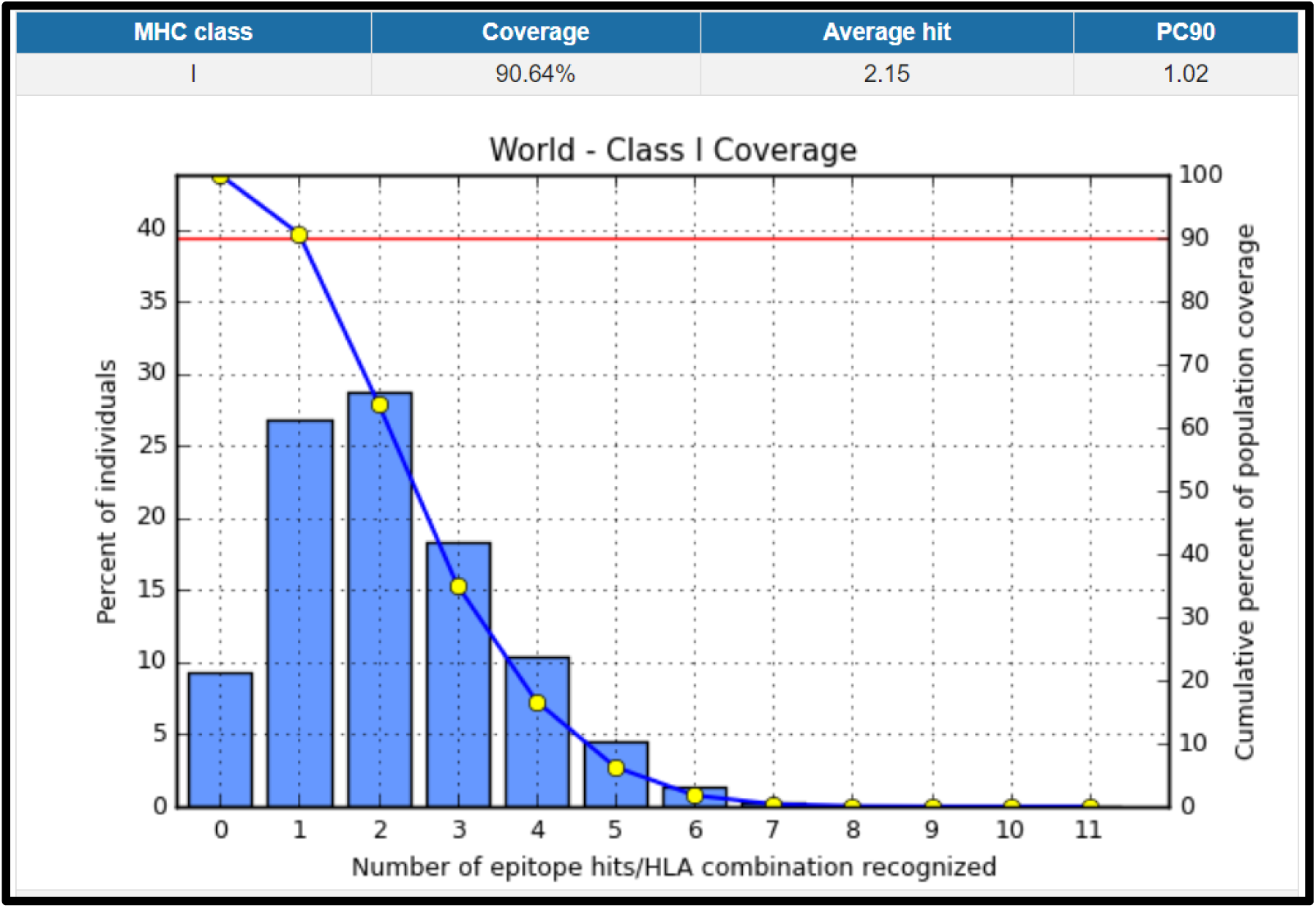
Illustrates the global resident’s total proportion for the top three MHC-I peptides (**YVYDTRGKL, FYRQGAFEL,** and **FTQLVAAYL**). Notes: In the graphs, the line (-o-) represents the cumulative percentage of population coverage of the epitopes; the bars represent the population coverage for each epitope

**Figure 17.**
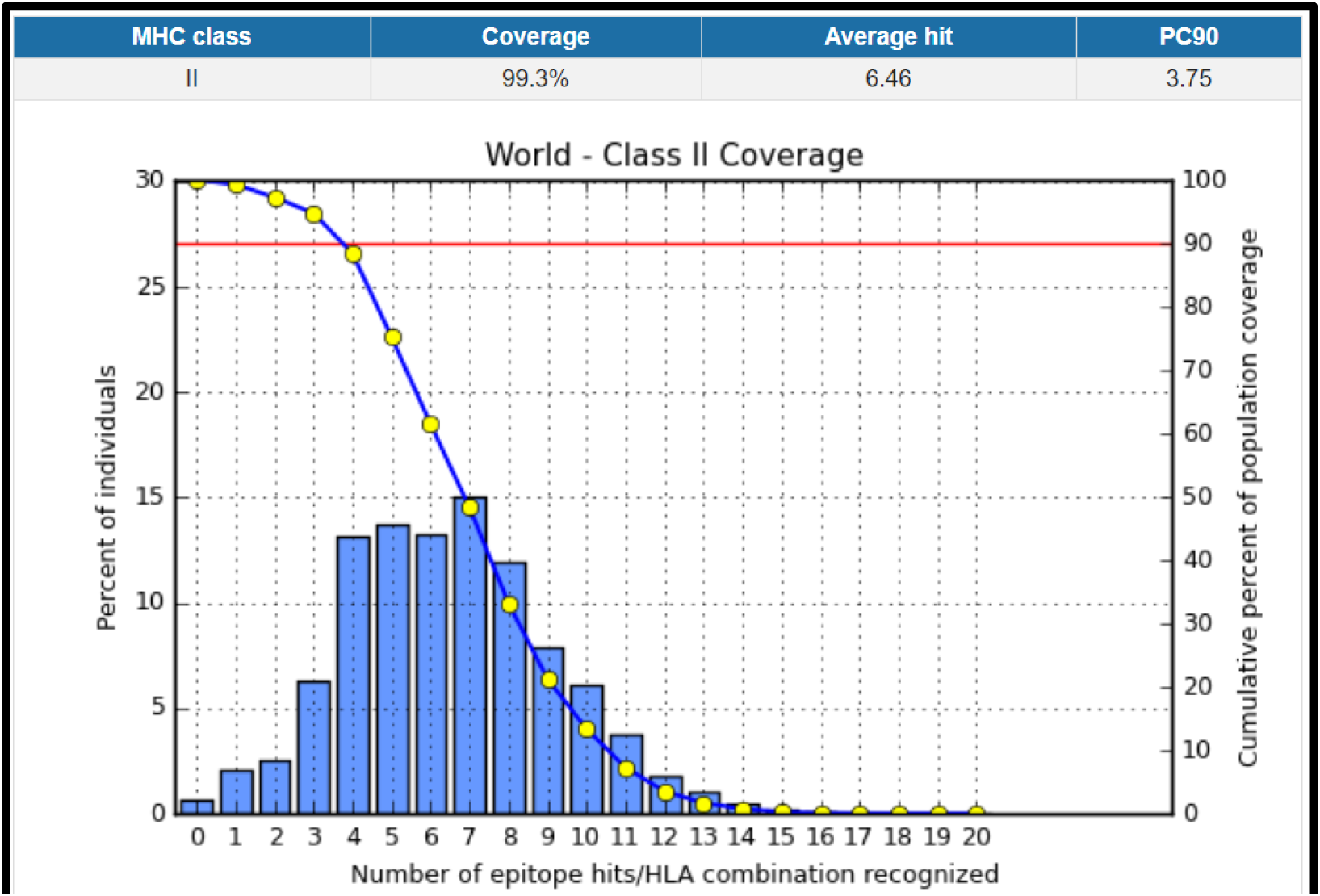
Illustrates the global resident’s total proportion for the top three MHC-II epitopes (**FFGGKVLNF, FYRQGAFEL, and FDYALVQHF**). Notes: In the graph, the line (-o-) represents the cumulative percentage of population coverage of the epitopes; the bars represent the population coverage for each epitope.

#### 3.4.2 Population coverage for MHC-I and MHC-II alleles combined

Regarding MHC-I and MHC-II alleles combined, three epitopes were supposed to interact with most predominant MHC class I and MHC class II alleles (**FFGGKVLNF, FYRQGAFEL,** and **FINAQLVDV**); represent a significant global coverage by IEDB population coverage tool. The most abundant population coverage percentage of these epitopes in the World was granted to **FFGGKVLNF** with percentage of 98.02%. The remarkable finding in this test is the average for the most abundant binders to combined MHC-I and MHC-II alleles; reveal an outstanding coverage with percentage of 99.91%.

**Table 6.** Global population coverage of the ten promising epitopes in MHC class I and II combined mode.

| Core epitope | Coverage<br>World class<br>I&II Combined | Total HLA<br>hits |
| --- | --- | --- |
| **FFGGKVLNF\*** | 98.20% | 12 |
| **FYRQGAFEL\*** | 97.95% | 16 |
| **FINAQLVDV\*** | 96.76% | 10 |
| FDYALVQHF | 96.32% | 14 |
| LVQHFAEEF | 95.27% | 8 |
| FSFTQLVAA | 92.03% | 8 |
| FACASLSPV | 87.11% | 11 |
| FTQLVAAYL | 82.74% | 15 |
| AAAALREAL | 79.45% | 6 |
| LEAAYADPA | 76.76% | 6 |
| YVYDTRGKL | 71.62% | 12 |
| **epitope set** | **88.56%** |  |

\*Top promising epitopes with massive population coverage

**Figure 18.**
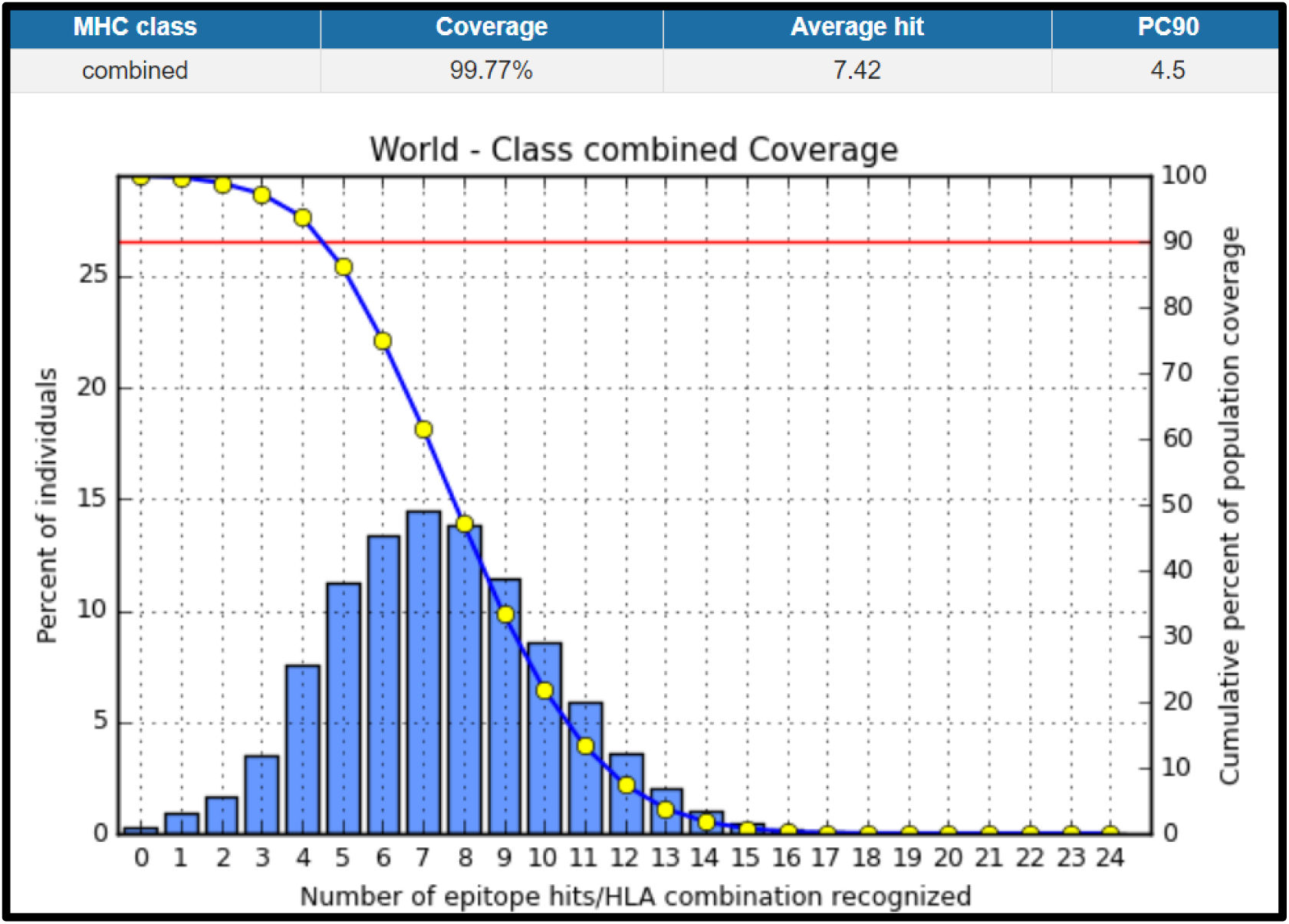
Illustrates the global population proportion for the top three MHC-I & II epitopes in combined mode (**FFGGKVLNF, FYRQGAFEL,** and **FINAQLVDV**). Notes: In the graphs, the line (-o-) represents the cumulative percentage of population coverage of the epitopes; the bars represent the population coverage for each epitope.

### 3.5 Molecular docking analysis

The 3D and 2D interactions of the models that had the strongest binding affinity among each epitope were represented in figures 19-25.

**Figure 19.**
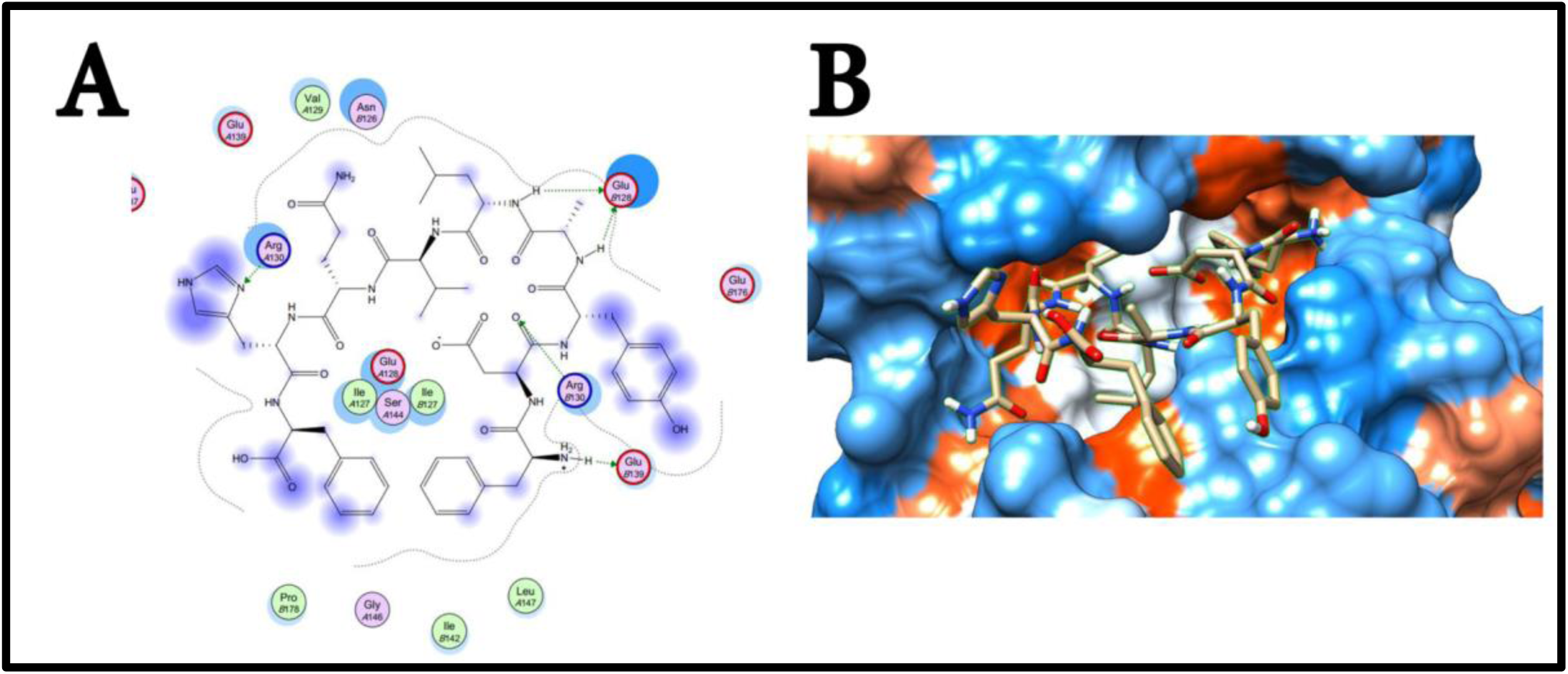
Illustrate the 3D interaction of best docking pose of **FDYALVQHF** in the binding sites of HLA-DRB1*01:01 using UCSF chimera 1.13.1 software (B), and 2D interaction (A) using Moe 2007.

**Figure 20.**
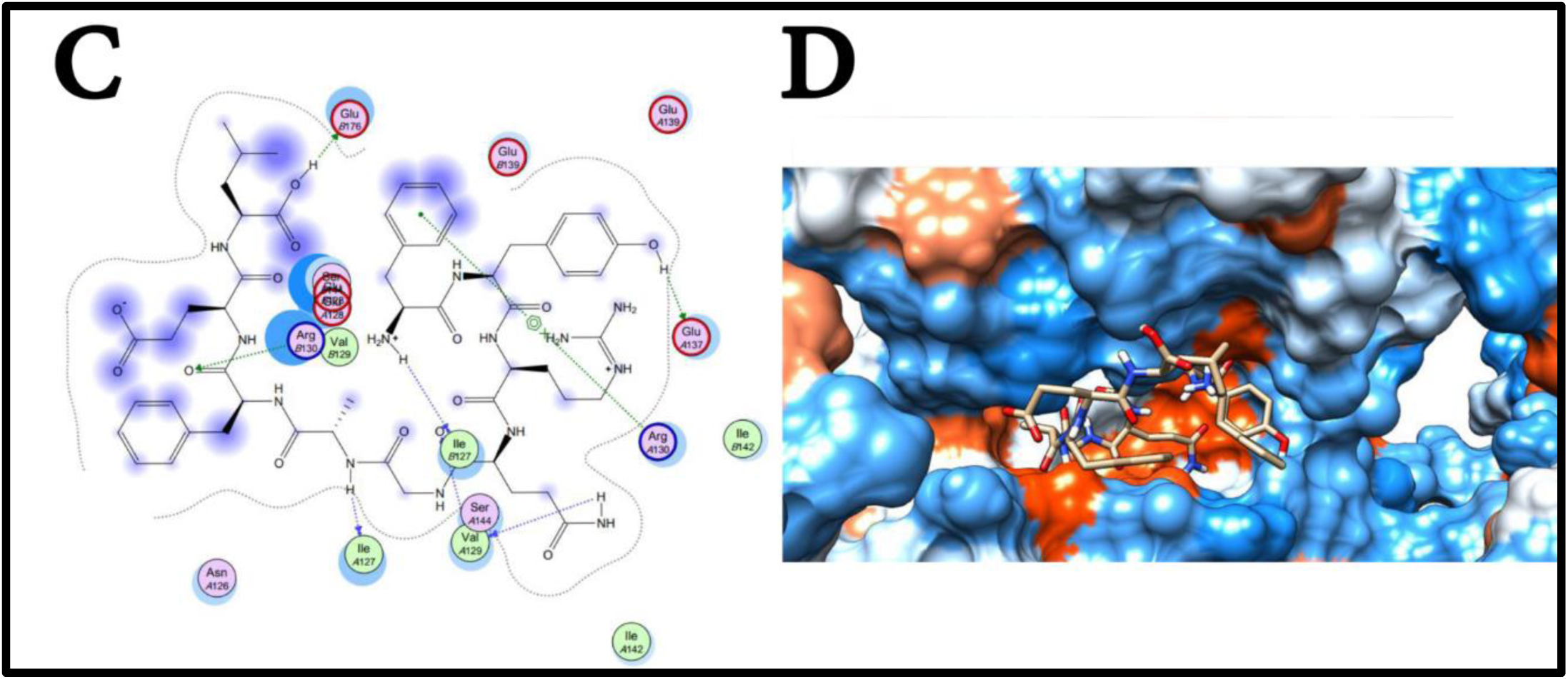
Illustrate the 3D interaction of best docking pose of **FYRQGAFEL** in the binding sites of HLA-DRB1*01:01 using UCSF chimera 1.13.1 software (D), and 2D interaction(C) using Moe 2007.

**Figure 21.**
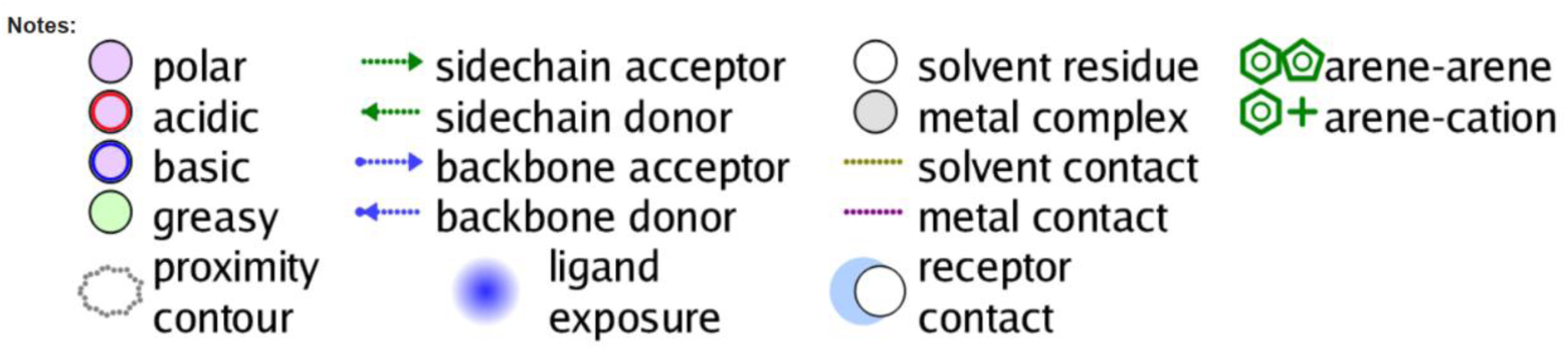
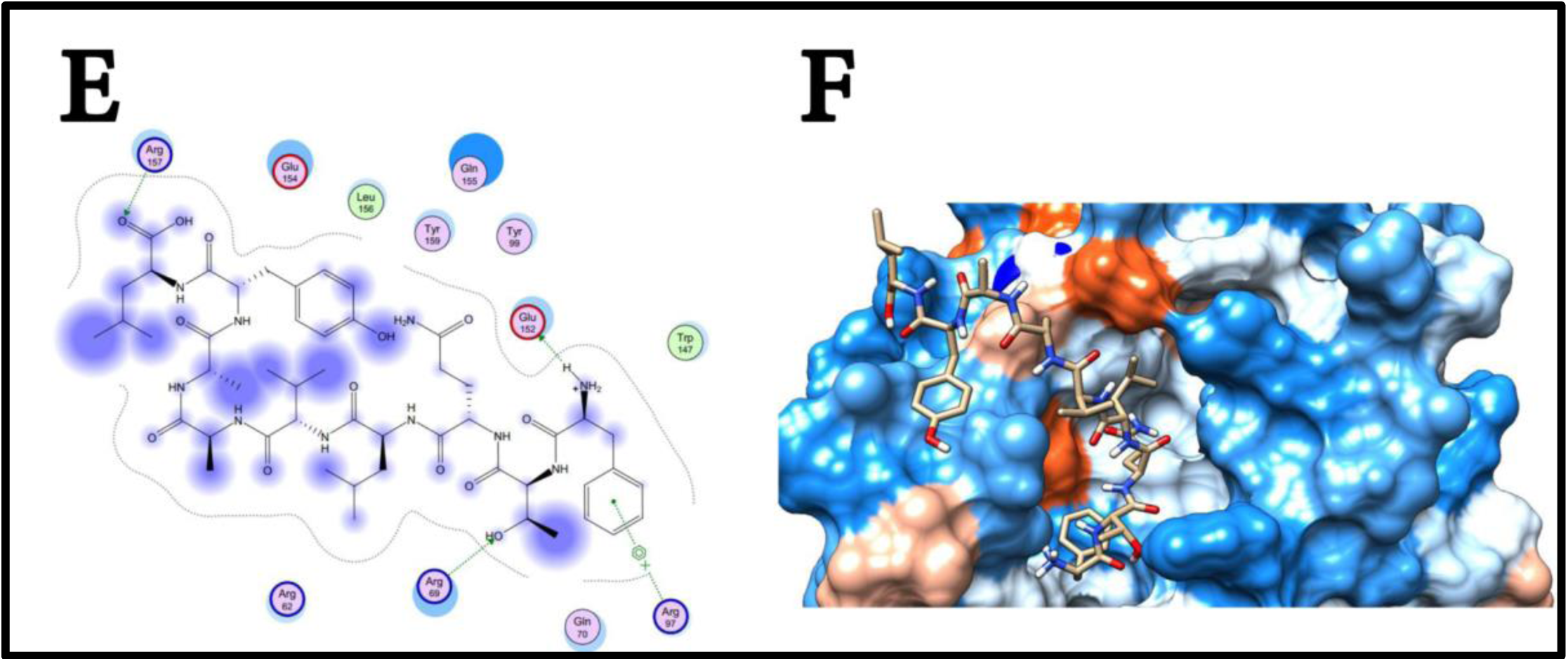
Illustrate the 3D interaction of best docking pose of **FTQLVAAYL** in the binding sites of HLA-C*12:03 using UCSF chimera 1.13.1 software (F), and 2D interaction (E) using Moe 2007.

**Figure 22.**
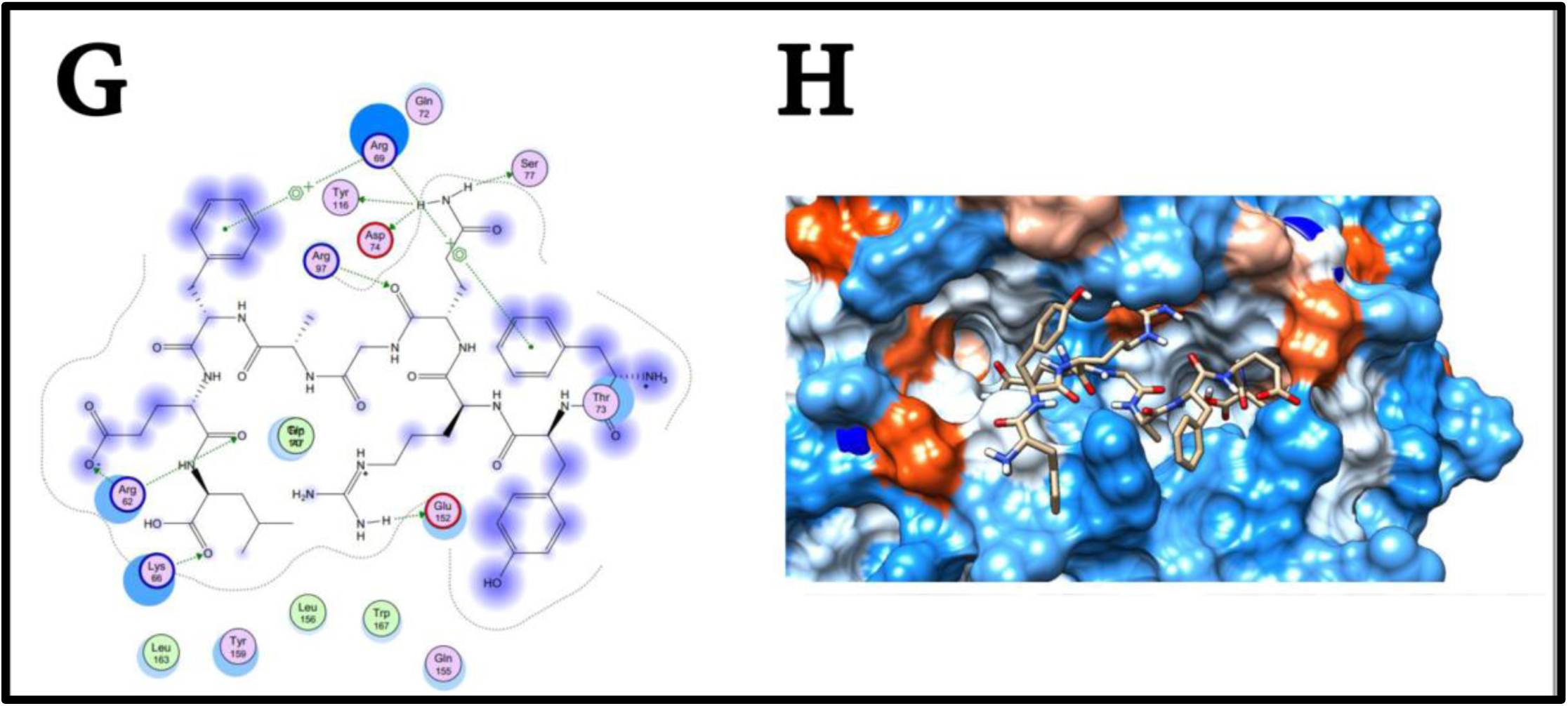
Illustrate the 3D interaction of best docking pose of **FYRQGAFEL** in the binding sites of HLA-C*12:03 using UCSF chimera 1.13.1 software (H), and 2D interaction (G) using Moe 2007.

**Figure 23.**
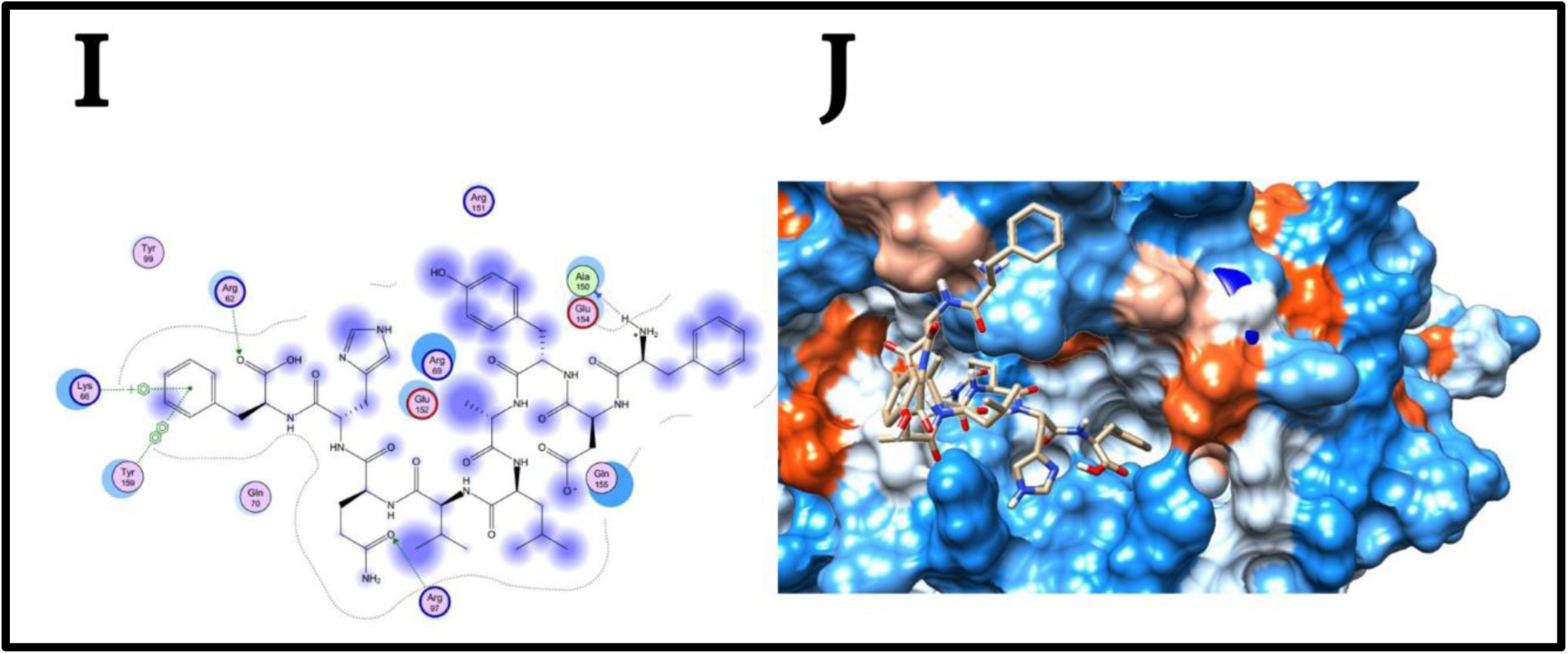
Illustrate the 3D interaction of best docking pose of **FDYALVQHF** in the binding sites of HLA-C*12:03 using UCSF chimera 1.13.1 software (I), and 2D interaction (J) using Moe 2007.

**Figure 24.**
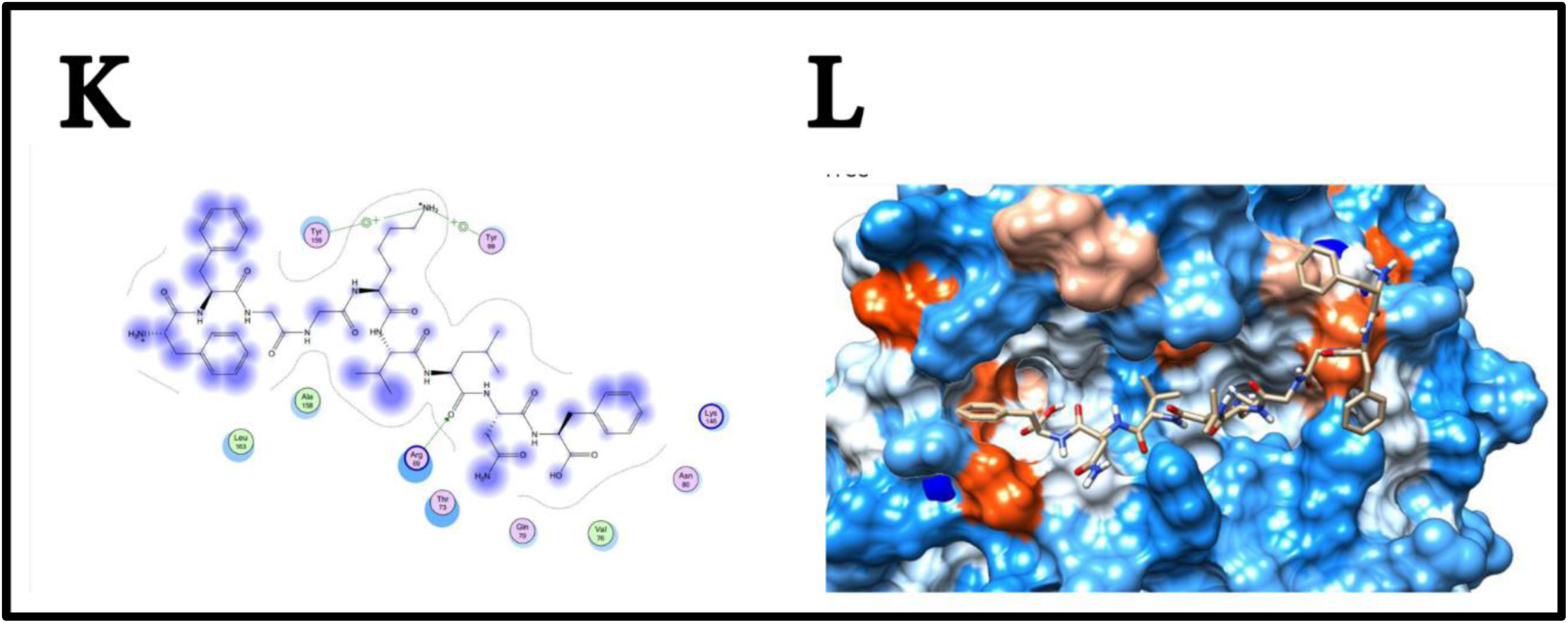
Illustrate the 3D interaction of best docking pose of **FFGGKVLNF** in the binding sites of HLA-C*12:03 using UCSF chimera 1.13.1 software (L), and 2D interaction (K) using Moe 2007.

**Figure 25.**
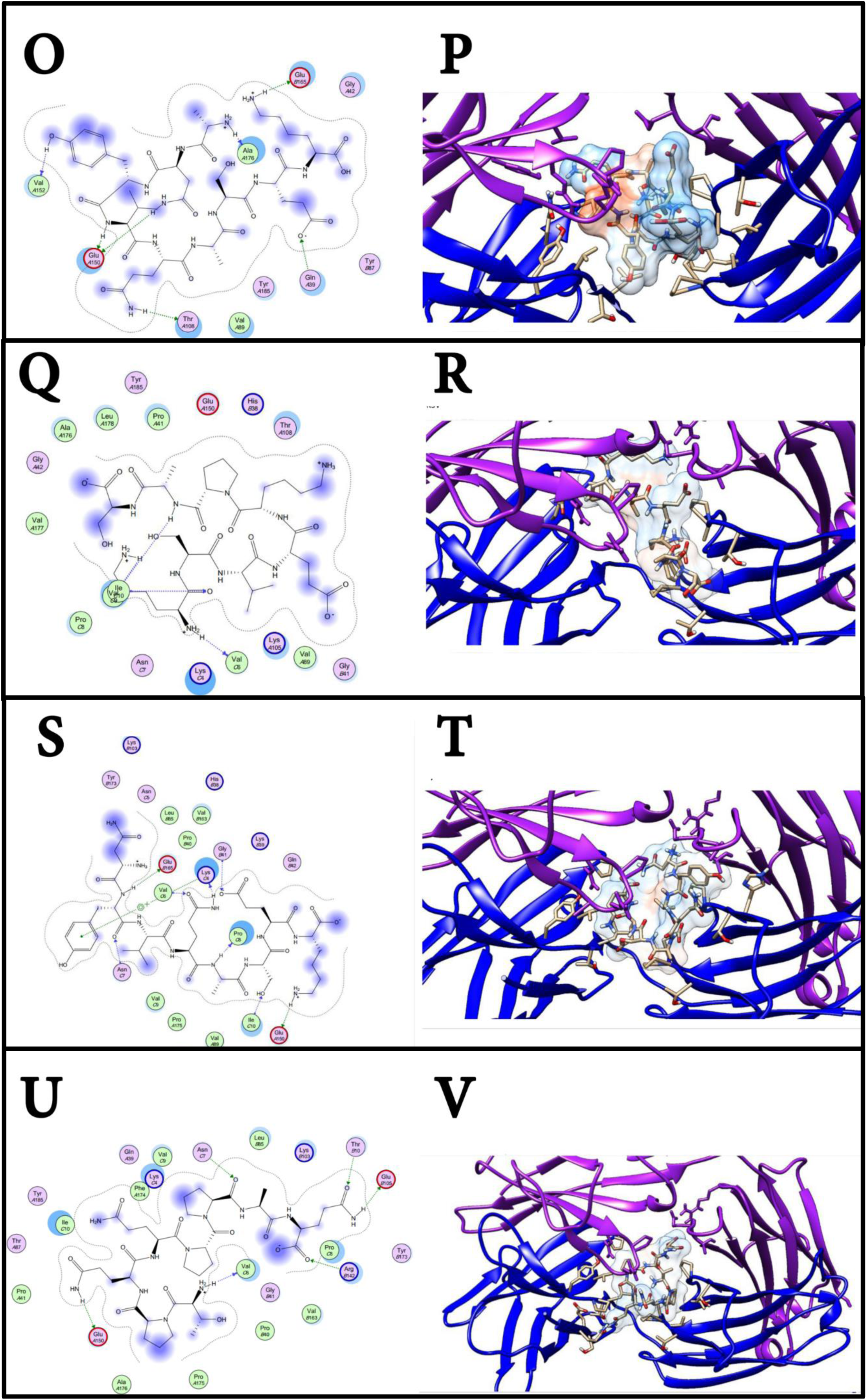
Illustrate the 3D and 2D interaction of the best docking poses of **ANYVQASEK, KSVEKPAS, NYVQASEK,** and **TPQQPPAQ** respectively, in the binding sites of Immunoglobulin G.

## 4. Discussion

The overall analysis revealed thirteen promiscuous B-cell and T-cell epitopes that consist of four immunogenic continuous B-cell epitopes (**ANYVQASEK, NYVQASEK, KSVEKPAS,** and **TPQQPPAQ)**, seven discontinuous B-cell epitopes, three immunogenic MHC-I epitopes (**YVYDTRGKL, FYRQGAFEL,** and **FTQLVAAYL)**, and three immunogenic MHC-II epitopes (**FFGGKVLNF, FYRQGAFEL,** and **FDYALVQHF)**, that proposed to be used in multi-epitope peptide vaccine designing.

Molecular docking and population coverage analysis are crucial factor in the development and refinement of epitopes selection. The epitope **FYRQGAFEL** had exhibited an exceptional result in terms of its broad spectrum of binding with MHC-I, MHC-II, and population coverage percentage. With universal coverage of 55.50%, 95.39% in MHC-I, MHC-II respectively, and 97.95% in combined mode. This infer a massive global population coverage, alongside it displayed the strongest binding affinity, when was docked with HLA-C*12:03 (ΔG −28.8 kcal/mol) over **FTQLVAAY** and **YVYDTRGKL**. and ΔG value of −37.35 kcal/mol when was docked with HLA-DRB1*01:01. Despite this outstanding coverage and binding score, **FYRQGAFEL** was not recognized as one of the abundant binders to MHC-I, a possible explanation that for its binding to one or more of the commonly occurring MHC-I alleles among global residents. The epitope **FDYALVQHF** also showed more negative free energy of binding (−46.25 kcal/mol) with HLA-DRB1*01:01 than **FYRQGAFEL**, which revealed a favored stability of the **FDYALVQHF** –MHC-II complex. Beside that **FDYALVQHF** had the abundant binding profile to MHC-II alleles and had the top population coverage of 95.38%. The core epitope **FFGGKVLNF** had the most dominant population coverage of 98.02% in MHC-II and 98.20% in combined mode. Even though **FFGGKVLNF** has the highest coverage among **FYRQGAFEL** and **FDYALVQHF**, it shows the weakest binding affinity (−31.05 kcal/mol) to HLA-DRB1*01:01 among them. **FFGGKVLNF** is considered a good candidate despite its weak binding affinity. Regarding MHC-I binding and population coverage, our finding has shown **YVYDTRGKL** that had the utmost binder and coverage of 60.93% with the highest global energy of −24.32 kcal/mol among **FYRQGAFEL** and **FTQLVAAY.** When it comes to the promising B-cell epitopes, **ANYVQASEK** and **TPQQPPAQ** had the strongest affinity toward Ig G with binding energy of −27.33 and -.54 kcal/mol respectively. **KSVEKPAS** and **NYVQASEK** come in the second place in terms of bind affinity. Apart from the ΔG binding value, the interaction between epitope and the chosen models can also be studied by analyzing the hydrogen bond between them. As in the case of the core peptide **FYRQGAFEL** showed eight hydrogen bonds were present in **FYRQGAFEL**–HLA-DRB1*01:01 complex and ten in **FYRQGAFEL**– HLA-C*12:03 complex. This might contribute the difference in binding energy.

Multi-epitope peptide-based vaccines are showing promising results. This emerging technology has facilitated the prevention and treatment of cancer, viral, bacterial and other diseases (95–100). This and other works were developed peptide vaccine against Salmonella, cholera, Mycobacterium and many other; imply the progressing of computational vaccinology approach (127, 181–201). However, this field is still in its infancy and there is dire need for further wet laboratory study. It is a widely held view the importance of Hsp70 family proteins as stand-alone virulence factors and immune response modulators since it prolongs the survival rate of mice against pulmonary Cryptococcal infection (7, 46, 66, 68–75). Several evidence of lines have suggested that many of heat shock proteins family as a potential candidate in designing of recombinant vaccine in mice models; Hsp 90 in *candida*, Hsp 60 in *Histoplasma* and Hsp 70 in *Schistosoma* (46, 66, 85–87). A study conducted in Hsp 70 of *Trypanosoma Cruzi* found four immunodominant epitopes (**TLLTIDGGI, DSLTNLRAL**,**TLQPVERVL** and **RIPKVMQLV**) were assayed for their recognition by CTL of HLA-A*02:01 and *T. cruzi*-infected transgenic B6-A2/Kb mice, two of them (**TLQPVERVL** and **RIPKVMQLV**) were also recognized by CTL of HLA-A*02:01 Chagas disease patients, indicating that these peptides are processed and displayed as MHC-I epitopes during the natural history of *T. cruzi* infection (82). An immunuinformatic study in immunoreactive mannoprotein MP88 of *Cryptococcus neoformans var. grubii* found three potenial MHC-I, MHC-II epitopes for each (**YMAADQFCL, VSYEEWMNY,** and **FQQRYTGTF)**, (**YARLLSLNA, ISYGTAMAV** and **INQTSYARL)** correspondingly, and four promising B-cell epitopes (**AYSTPA, AYSTPAS, PASSNCK,** and **DSAYPP)** (202). Thus, this support our findings and point toward to the fact that the development of a Cryptococcal vaccine is feasible and possible through screening the *Cryptococcus neoformans*’s immunogenic proteins and utilize the promising antigenic epitopes in peptide vaccine designing. A therapeutic vaccine that able to prevent reactivation and be effective in the setting of established Cryptococcosis (29, 62). There are experiments with conventional vaccines in the *C. neoformans* field, killed vaccines have generally been ineffective and some have enhanced infection. Live vaccines using attenuated mutants have been shown to induce stronger, longer-lasting immune responses in immunocompetent (203–205). However, live vaccines are not safe for use in immunocompromised patients and any attempt to develop a live vaccine for cryptococcosis is likely to face significant ethical outcome. In contrast, the success of subunit and conjugate vaccines against Hepatitis B virus, *Haemophilus influenza* type B and *Streptococcus pneumonia* has shown the safety and effectiveness of this approach (206). In patients with suppressed T-cell responses will undoubtedly suffer from reduced memory responses, rendering conventional vaccine strategies useless. Hence, implementing novel combined T-cell and B-cell vaccines that have the potential to mediate protective immunity against *C. neoformans* would improve the quality of life of immunocompromised patients (26). Nonetheless, the efficacy of Cryptococcus vaccine candidate to induce protection against cryptococcosis will need to be confirmed using an immune-deficient animal model system to mimic immune suppression in human populations (76).

The findings in this study are subjected to few limitations, a limited number of validated sequences were retrieved might due to the lack of equivalent data in the literature; biases could be incorporated. Furthermore, regarding HLA allele frequencies and reference sets with population coverage, there is no predictor for (HLA-DRB5*01:01, HLA-DPA1*01 and HLA-DRB3*01:01) at IEDB population coverage tool, which might mislead the inference of coverage percentage.

On the other hand, computational vaccinology approach speeds up the process of successful identification of potential peptide-vaccine candidates and greatly downsize the number of epitopes to be synthesized and analyzed for experimental assays. Therefore, there is a definite need for experimental validation for the carefully chosen vaccine candidates *in vitro* and *in vivo* to fortify their antigenic and immunogenic potentials. Additionally, further computational studies are needed to be conducted in pathogens that expressed Hsp70, as it believed to find out universal epitopes that might be overlapped with other pathogens-derived Hsp70. Finally, *C. neoformans* expresses a significant number of virulence agents that could help the parasite to evade and evoke host immunity. Thus, screening of new immune-proteomic factors may facilitate the future development of therapeutic interventions aimed at boosting human being immunity against cryptococcosis.

Theoretically, No single epitope vaccine would provide a universal protection against all *Cryptococcus neoformans var. grubii* strains because of allelic polymorphism among global population, and epitopes have a different binding profile with different HLA alleles (59, 157). Nevertheless, a complete protection would be achieved by combining multiple epitopes by targeting immunodominant regions comprising of multiple epitopes. Thus, our prime vaccine candidate was a putative ten promising epitopes (**ANYVQASEK, NYVQASEK, KSVEKPAS, TPQQPPAQ, YVYDTRGKL, FYRQGAFEL, FTQLVAAYL, FFGGKVLNF, FDYALVQHF,** and **FINAQLVDV**). Together, these epitopes are forecasted to trigger T lymphocytes, B lymphocytes, and immunological memory with overall coverage above 90%. Accordingly, our *in silico* vaccine is expected to be the future multi-epitope peptide vaccine with potential immunogenicity and minimum allergenicity that able to stimulate desirable immune responses against all strain of *Cryptococcus neoformans var. grubii* with massive global population coverage.

## 5. Conclusion

Cryptococcosis is a serious global problem concerning morbidity and mortality in immunocompromised individuals. Unfortunately, the unavailability of vaccines and the failure of antifungal against Cryptococcosis has led to affect many precious lives in various regions of the world. However, Structural and genomic data, alongside the drastic development of vast genome sequence databases, aids in the design and discovery of novel vaccine candidates when coupled together with computational tools. We inferred that the predicted epitopes possess therapeutic potential, with promising scope in the near future. Our *in silico* analysis provide novel insights regarding computational vaccinology, which will aid in the development of potential peptide vaccines using the predicted peptides.

