## Supplementary material for "Computational vaccinology approach: Designing an efficient multi-epitope peptide vaccine against *Cryptococcus neoformans var. grubii’s* heat shock 70KDa protein": supplemantal Table 2

### Supplementary Tables and Figures

**Table 2.** List of promising epitopes that had a good binding affinity with MHC-I alleles in terms of $IC_{50}$ and Percentile rank.

| epitopes | Start | End | Allele | $IC_{50}$ | Percentile |
|---|---|---|---|---|---|
| **YVYDTRGKL*** | 579 | 587 | HLA-A*02:06 | 274.73 | 1.8 |
| | 579 | 587 | HLA-A*68:02 | 359.64 | 1.5 |
| | 579 | 587 | HLA-B*07:02 | 488.64 | 1.3 |
| | 579 | 587 | HLA-C*03:03 | 10.15 | 0.06 |
| | 579 | 587 | HLA-C*06:02 | 350.13 | 0.13 |
| | 579 | 587 | HLA-C*07:01 | 133.84 | 0.04 |
| | 579 | 587 | HLA-C*12:03 | 10.95 | 0.03 |
| | 579 | 587 | HLA-C*14:02 | 10.45 | 0.02 |
| | 579 | 587 | HLA-C*15:02 | 475.11 | 0.24 |
| **LTFYRQGAF*** | 435 | 443 | HLA-A*29:02 | 497.2 | 1.5 |
| | 435 | 443 | HLA-A*32:01 | 198.13 | 0.23 |
| | 435 | 443 | HLA-B*14:02 | 435.58 | 0.03 |
| | 435 | 443 | HLA-B*15:01 | 34.93 | 0.21 |
| | 435 | 443 | HLA-C*03:03 | 427.85 | 0.68 |
| | 435 | 443 | HLA-C*12:03 | 478.63 | 0.53 |
| | 435 | 443 | HLA-C*14:02 | 464.12 | 0.54 |
| **RATPSLVSF*** | 34 | 42 | HLA-A*02:06 | 480.08 | 2.5 |
| | 34 | 42 | HLA-B*15:01 | 153.24 | 0.73 |
| | 34 | 42 | HLA-B*35:01 | 346.58 | 0.65 |
| | 34 | 42 | HLA-B*57:01 | 79.07 | 0.31 |
| | 34 | 42 | HLA-B*58:01 | 15.06 | 0.08 |
| | 34 | 42 | HLA-C*03:03 | 19 | 0.11 |
| | 34 | 42 | HLA-C*12:03 | 70.21 | 0.15 |
| **FTQLVAAYL*** | 115 | 123 | HLA-A*02:01 | 387.43 | 2.6 |
| | 115 | 123 | HLA-A*02:06 | 166.27 | 1.4 |
| | 115 | 123 | HLA-A*68:02 | 21.89 | 0.22 |
| | 115 | 123 | HLA-C*03:03 | 335.75 | 0.6 |
| | 115 | 123 | HLA-C*05:01 | 356.66 | 0.18 |
| | 115 | 123 | HLA-C*14:02 | 74.62 | 0.14 |
| | 115 | 123 | HLA-C*15:02 | 276.33 | 0.16 |
| FVDVGHSDY | 201 | 209 | HLA-A*01:01 | 7.75 | 0.03 |
| | 201 | 209 | HLA-A*29:02 | 21.21 | 0.16 |
| | 201 | 209 | HLA-B*15:02 | 249.36 | 0.1 |
| | 201 | 209 | HLA-B*35:01 | 39.17 | 0.15 |
| | 201 | 209 | HLA-C*05:01 | 48.13 | 0.06 |
| FACASLSPV | 379 | 387 | HLA-A*02:01 | 73.02 | 0.74 |
| | 379 | 387 | HLA-A*02:06 | 4.38 | 0.03 |
| | 379 | 387 | HLA-A*68:02 | 16.12 | 0.16 |

| | | | | | |
|---|---|---|---|---|---|
| | 379 | 387 | HLA-C*03:03 | 17.16 | 0.1 |
| | 379 | 387 | HLA-C*12:03 | 11.68 | 0.03 |
| | 379 | 387 | HLA-C*15:02 | 499.4 | 0.25 |
| YADPASLPK | 449 | 457 | HLA-A*11:01 | 101.43 | 0.69 |
| | 449 | 457 | HLA-A*68:01 | 391.98 | 1.9 |
| | 449 | 457 | HLA-B*35:01 | 470.2 | 0.8 |
| | 449 | 457 | HLA-C*03:03 | 8.24 | 0.05 |
| | 449 | 457 | HLA-C*05:01 | 77.92 | 0.08 |
| | 449 | 457 | HLA-C*12:03 | 123.82 | 0.21 |
| GIMNFEGAY | 492 | 500 | HLA-A*11:01 | 262.9 | 1.6 |
| | 492 | 500 | HLA-A*29:02 | 147.9 | 0.68 |
| | 492 | 500 | HLA-A*30:02 | 28.97 | 0.06 |
| | 492 | 500 | HLA-B*15:01 | 91.16 | 0.49 |
| | 492 | 500 | HLA-B*15:02 | 282.81 | 0.11 |
| | 492 | 500 | HLA-B*35:01 | 468.92 | 0.8 |
| FYRQGAFEL | 437 | 445 | HLA-A*23:01 | 163.92 | 0.47 |
| | 437 | 445 | HLA-A*24:02 | 379.48 | 0.6 |
| | 437 | 445 | HLA-C*03:03 | 234.95 | 0.5 |
| | 437 | 445 | HLA-C*07:02 | 27.33 | 0.02 |
| | 437 | 445 | HLA-C*12:03 | 429.65 | 0.5 |
| | 437 | 445 | HLA-C*14:02 | 10.8 | 0.02 |
| GAFELEAAY | 441 | 449 | HLA-A*29:02 | 152.13 | 0.69 |
| | 441 | 449 | HLA-A*30:02 | 204.96 | 0.78 |
| | 441 | 449 | HLA-B*15:01 | 175.23 | 0.79 |
| | 441 | 449 | HLA-B*35:01 | 12.71 | 0.06 |
| | 441 | 449 | HLA-B*46:01 | 311.25 | 0.05 |
| | 441 | 449 | HLA-C*12:03 | 497.7 | 0.54 |

*****Top promising epitopes with efficient binding affinity
