## Supplementary material for "Computational vaccinology approach: Designing an efficient multi-epitope peptide vaccine against *Cryptococcus neoformans var. grubii’s* heat shock 70KDa protein": supplemantal Table 3

### Supplementary Tables and Figures

**Table 3.** List of the promising epitopes core sequence that had an efficient binding affinity with MHC-II in terms of $IC_{50}$ and Percentile Ranks.

| Core Sequence | Start | End | Allele | Epitope Sequence | IC50 | Rank |
|---|---|---|---|---|---|---|
| **FDYALVQHF*** | 234 | 248 | HLA-DPA1*01:03/DPB1*02:01 | GRDFDYALVQHFAEE | 86.1 | 7.57 |
| | 233 | 247 | HLA-DPA1*01:03/DPB1*02:01 | GGRDFDYALVQHFAE | 94.9 | 8.08 |
| | 235 | 249 | HLA-DPA1*02:01/DPB1*01:01 | RDFDYALVQHFAEEF | 40.4 | 4.14 |
| | 234 | 248 | HLA-DPA1*02:01/DPB1*01:01 | GRDFDYALVQHFAEE | 45.6 | 4.76 |
| | 233 | 247 | HLA-DPA1*02:01/DPB1*01:01 | GGRDFDYALVQHFAE | 56.2 | 5.99 |
| | 232 | 246 | HLA-DPA1*02:01/DPB1*01:01 | FGGRDFDYALVQHFA | 67.3 | 7.21 |
| | 236 | 250 | HLA-DPA1*02:01/DPB1*01:01 | DFDYALVQHFAEEFK | 67.4 | 7.22 |
| | 231 | 245 | HLA-DPA1*02:01/DPB1*01:01 | HFGGRDFDYALVQHF | 78.6 | 8.4 |
| | 234 | 248 | HLA-DRB1*01:01 | GRDFDYALVQHFAEE | 54.3 | 22.25 |
| | 233 | 247 | HLA-DRB1*01:01 | GGRDFDYALVQHFAE | 66.6 | 24.79 |
| | 235 | 249 | HLA-DRB1*01:01 | RDFDYALVQHFAEEF | 70.6 | 25.53 |
| | 232 | 246 | HLA-DRB1*01:01 | FGGRDFDYALVQHFA | 83.7 | 27.76 |
| | 233 | 247 | HLA-DRB1*03:01 | GGRDFDYALVQHFAE | 28.2 | 1.65 |
| | 234 | 248 | HLA-DRB1*03:01 | GRDFDYALVQHFAEE | 28.2 | 1.65 |
| | 232 | 246 | HLA-DRB1*03:01 | FGGRDFDYALVQHFA | 32 | 1.88 |
| | 231 | 245 | HLA-DRB1*03:01 | HFGGRDFDYALVQHF | 45 | 2.63 |
| | 235 | 249 | HLA-DRB1*03:01 | RDFDYALVQHFAEEF | 66.6 | 3.64 |
| | 234 | 248 | HLA-DRB1*04:05 | GRDFDYALVQHFAEE | 21.1 | 1.41 |
| | 233 | 247 | HLA-DRB1*04:05 | GGRDFDYALVQHFAE | 23.7 | 1.73 |
| | 235 | 249 | HLA-DRB1*04:05 | RDFDYALVQHFAEEF | 23.7 | 1.73 |
| | 236 | 250 | HLA-DRB1*04:05 | DFDYALVQHFAEEFK | 30.6 | 2.55 |
| | 232 | 246 | HLA-DRB1*04:05 | FGGRDFDYALVQHFA | 31.5 | 2.67 |
| | 231 | 245 | HLA-DRB1*04:05 | HFGGRDFDYALVQHF | 37.3 | 3.4 |
| | 237 | 251 | HLA-DRB1*04:05 | FDYALVQHFAEEFKT | 40.9 | 3.82 |
| | 232 | 246 | HLA-DRB1*07:01 | FGGRDFDYALVQHFA | 10.7 | 1.8 |
| | 231 | 245 | HLA-DRB1*07:01 | HFGGRDFDYALVQHF | 10.8 | 1.83 |
| | 233 | 247 | HLA-DRB1*07:01 | GGRDFDYALVQHFAE | 13 | 2.34 |
| | 235 | 249 | HLA-DRB1*07:01 | RDFDYALVQHFAEEF | 13.4 | 2.44 |
| | 234 | 248 | HLA-DRB1*07:01 | GRDFDYALVQHFAEE | 17 | 3.22 |
| | 236 | 250 | HLA-DRB1*07:01 | DFDYALVQHFAEEFK | 20.2 | 3.88 |
| | 237 | 251 | HLA-DRB1*07:01 | FDYALVQHFAEEFKT | 25.9 | 5 |
| | 235 | 249 | HLA-DRB1*09:01 | RDFDYALVQHFAEEF | 81.3 | 5.61 |
| | 234 | 248 | HLA-DRB1*11:01 | GRDFDYALVQHFAEE | 65.9 | 10.43 |
| | 234 | 248 | HLA-DRB5*01:01 | GRDFDYALVQHFAEE | 70.8 | 12.49 |
| | 233 | 247 | HLA-DRB5*01:01 | GGRDFDYALVQHFAE | 73.2 | 12.75 |
| | 232 | 246 | HLA-DRB5*01:01 | FGGRDFDYALVQHFA | 77.1 | 13.16 |
| | 235 | 249 | HLA-DRB5*01:01 | RDFDYALVQHFAEEF | 96.2 | 14.96 |
| | 231 | 245 | HLA-DRB5*01:01 | HFGGRDFDYALVQHF | 99.4 | 15.25 |

| | | | | | | |
|---|---|---|---|---|---|---|
| **FFGGKVLNF*** | 353 | 367 | HLA-DPA1*01:03/DPB1*02:01 | RIQQFFGGKVLNFTL | 57.8 | 5.71 |
| | 352 | 366 | HLA-DPA1*01:03/DPB1*02:01 | ERIQQFFGGKVLNFT | 65.4 | 6.25 |
| | 351 | 365 | HLA-DPA1*01:03/DPB1*02:01 | KERIQQFFGGKVLNF | 69.3 | 6.5 |
| | 354 | 368 | HLA-DPA1*01:03/DPB1*02:01 | IQQFFGGKVLNFTLN | 71.2 | 6.63 |
| | 354 | 368 | HLA-DPA1*02:01/DPB1*01:01 | IQQFFGGKVLNFTLN | 43.6 | 4.53 |
| | 355 | 369 | HLA-DPA1*02:01/DPB1*01:01 | QQFFGGKVLNFTLNQ | 71.2 | 7.63 |
| | 355 | 369 | HLA-DPA1*03:01/DPB1*04:02 | QQFFGGKVLNFTLNQ | 72.6 | 7.73 |
| | 352 | 366 | HLA-DQA1*05:01/DQB1*03:01 | ERIQQFFGGKVLNFT | 14.4 | 2.46 |
| | 353 | 367 | HLA-DQA1*05:01/DQB1*03:01 | RIQQFFGGKVLNFTL | 15 | 2.6 |
| | 354 | 368 | HLA-DQA1*05:01/DQB1*03:01 | IQQFFGGKVLNFTLN | 15.4 | 2.68 |
| | 355 | 369 | HLA-DQA1*05:01/DQB1*03:01 | QQFFGGKVLNFTLNQ | 18.4 | 3.36 |
| | 351 | 365 | HLA-DQA1*05:01/DQB1*03:01 | KERIQQFFGGKVLNF | 19.7 | 3.63 |
| | 356 | 370 | HLA-DQA1*05:01/DQB1*03:01 | QFFGGKVLNFTLNQD | 84.3 | 13.24 |
| | 353 | 367 | HLA-DRB1*01:01 | RIQQFFGGKVLNFTL | 20.4 | 11.5 |
| | 354 | 368 | HLA-DRB1*01:01 | IQQFFGGKVLNFTLN | 23.3 | 12.82 |
| | 355 | 369 | HLA-DRB1*01:01 | QQFFGGKVLNFTLNQ | 51.9 | 21.7 |
| | 353 | 367 | HLA-DRB1*07:01 | RIQQFFGGKVLNFTL | 29.2 | 5.56 |
| | 354 | 368 | HLA-DRB1*07:01 | IQQFFGGKVLNFTLN | 49.8 | 8.69 |
| **FTQLVAAYL*** | 112 | 126 | HLA-DRB1*01:01 | DFSFTQLVAAYLGKL | 4.3 | 0.25 |
| | 111 | 125 | HLA-DRB1*01:01 | TDFSFTQLVAAYLGK | 4.6 | 0.47 |
| | 113 | 127 | HLA-DRB1*01:01 | FSFTQLVAAYLGKLR | 4.6 | 0.47 |
| | 110 | 124 | HLA-DRB1*01:01 | PTDFSFTQLVAAYLG | 4.9 | 0.71 |
| | 109 | 123 | HLA-DRB1*01:01 | EPTDFSFTQLVAAYL | 5.2 | 0.97 |
| | 114 | 128 | HLA-DRB1*01:01 | SFTQLVAAYLGKLRD | 5.5 | 1.24 |
| | 115 | 129 | HLA-DRB1*01:01 | FTQLVAAYLGKLRDT | 6.7 | 2.37 |
| | 113 | 127 | HLA-DRB1*04:04 | FSFTQLVAAYLGKLR | 25.9 | 2.24 |
| | 112 | 126 | HLA-DRB1*04:04 | DFSFTQLVAAYLGKL | 28.3 | 2.59 |
| | 111 | 125 | HLA-DRB1*04:04 | TDFSFTQLVAAYLGK | 30.1 | 2.83 |
| | 110 | 124 | HLA-DRB1*04:04 | PTDFSFTQLVAAYLG | 32.5 | 3.18 |
| | 109 | 123 | HLA-DRB1*04:04 | EPTDFSFTQLVAAYL | 37.6 | 3.93 |
| | 114 | 128 | HLA-DRB1*04:04 | SFTQLVAAYLGKLRD | 44.9 | 4.96 |
| | 115 | 129 | HLA-DRB1*04:04 | FTQLVAAYLGKLRDT | 91.2 | 10.94 |
| | 109 | 123 | HLA-DRB1*04:05 | EPTDFSFTQLVAAYL | 84.2 | 8.25 |
| | 110 | 124 | HLA-DRB1*04:05 | PTDFSFTQLVAAYLG | 97.5 | 9.42 |
| | 109 | 123 | HLA-DRB1*07:01 | EPTDFSFTQLVAAYL | 12.9 | 2.32 |
| | 110 | 124 | HLA-DRB1*07:01 | PTDFSFTQLVAAYLG | 21.1 | 4.07 |
| | 112 | 126 | HLA-DRB1*07:01 | DFSFTQLVAAYLGKL | 31 | 5.87 |
| | 111 | 125 | HLA-DRB1*07:01 | TDFSFTQLVAAYLGK | 32.2 | 6.07 |
| | 113 | 127 | HLA-DRB1*07:01 | FSFTQLVAAYLGKLR | 36.6 | 6.7 |
| | 114 | 128 | HLA-DRB1*07:01 | SFTQLVAAYLGKLRD | 52.5 | 9.06 |
| | 115 | 129 | HLA-DRB1*07:01 | FTQLVAAYLGKLRDT | 70.4 | 11.2 |
| | 112 | 126 | HLA-DRB1*09:01 | DFSFTQLVAAYLGKL | 20 | 0.79 |
| | 111 | 125 | HLA-DRB1*09:01 | TDFSFTQLVAAYLGK | 23.3 | 1.04 |
| | 113 | 127 | HLA-DRB1*09:01 | FSFTQLVAAYLGKLR | 23.4 | 1.04 |

| | | | | | | |
|---|---|---|---|---|---|---|
| | 110 | 124 | HLA-DRB1*09:01 | PTDFSFTQLVAAYLG | 23.6 | 1.05 |
| | 109 | 123 | HLA-DRB1*09:01 | EPTDFSFTQLVAAYL | 25.5 | 1.21 |
| | 114 | 128 | HLA-DRB1*09:01 | SFTQLVAAYLGKLRD | 27.5 | 1.37 |
| | 115 | 129 | HLA-DRB1*09:01 | FTQLVAAYLGKLRDT | 37 | 2.15 |
| | 112 | 126 | HLA-DRB1*11:01 | DFSFTQLVAAYLGKL | 78.9 | 11.81 |
| | 113 | 127 | HLA-DRB1*11:01 | FSFTQLVAAYLGKLR | 97.9 | 13.55 |
| | 114 | 128 | HLA-DRB1*15:01 | SFTQLVAAYLGKLRD | 33.3 | 3.14 |
| | 113 | 127 | HLA-DRB1*15:01 | FSFTQLVAAYLGKLR | 36.1 | 3.47 |
| | 112 | 126 | HLA-DRB1*15:01 | DFSFTQLVAAYLGKL | 50.1 | 5.1 |
| | 113 | 127 | HLA-DRB5*01:01 | FSFTQLVAAYLGKLR | 9.7 | 2.28 |
| | 112 | 126 | HLA-DRB5*01:01 | DFSFTQLVAAYLGKL | 10.6 | 2.54 |
| | 114 | 128 | HLA-DRB5*01:01 | SFTQLVAAYLGKLRD | 12.1 | 2.96 |
| | 115 | 129 | HLA-DRB5*01:01 | FTQLVAAYLGKLRDT | 15.7 | 3.89 |
| | 111 | 125 | HLA-DRB5*01:01 | TDFSFTQLVAAYLGK | 17.1 | 4.21 |
| | 110 | 124 | HLA-DRB5*01:01 | PTDFSFTQLVAAYLG | 29.9 | 6.84 |
| | 109 | 123 | HLA-DRB5*01:01 | EPTDFSFTQLVAAYL | 37 | 8.08 |
| **IAGLNALRL*** | 160 | 174 | HLA-DPA1*03:01/DPB1*04:02 | AANIAGLNALRLIND | 78.6 | 8.2 |
| | 159 | 173 | HLA-DPA1*03:01/DPB1*04:02 | DAANIAGLNALRLIN | 85.3 | 8.68 |
| | 159 | 173 | HLA-DRB1*01:01 | DAANIAGLNALRLIN | 4.8 | 0.62 |
| | 160 | 174 | HLA-DRB1*01:01 | AANIAGLNALRLIND | 4.8 | 0.62 |
| | 158 | 172 | HLA-DRB1*01:01 | LDAANIAGLNALRLI | 5.2 | 0.97 |
| | 161 | 175 | HLA-DRB1*01:01 | ANIAGLNALRLINDN | 5.6 | 1.33 |
| | 157 | 171 | HLA-DRB1*01:01 | LLDAANIAGLNALRL | 6.7 | 2.37 |
| | 162 | 176 | HLA-DRB1*01:01 | NIAGLNALRLINDNT | 7 | 2.64 |
| | 163 | 177 | HLA-DRB1*01:01 | IAGLNALRLINDNTA | 9 | 4.39 |
| | 157 | 171 | HLA-DRB1*07:01 | LLDAANIAGLNALRL | 20.9 | 4.03 |
| | 158 | 172 | HLA-DRB1*07:01 | LDAANIAGLNALRLI | 21.4 | 4.14 |
| | 159 | 173 | HLA-DRB1*07:01 | DAANIAGLNALRLIN | 38.1 | 6.93 |
| | 160 | 174 | HLA-DRB1*07:01 | AANIAGLNALRLIND | 81.2 | 12.34 |
| | 160 | 174 | HLA-DRB1*09:01 | AANIAGLNALRLIND | 25.3 | 1.2 |
| | 159 | 173 | HLA-DRB1*09:01 | DAANIAGLNALRLIN | 29.6 | 1.53 |
| | 158 | 172 | HLA-DRB1*09:01 | LDAANIAGLNALRLI | 32.7 | 1.78 |
| | 161 | 175 | HLA-DRB1*09:01 | ANIAGLNALRLINDN | 36.1 | 2.06 |
| | 157 | 171 | HLA-DRB1*09:01 | LLDAANIAGLNALRL | 38.3 | 2.26 |
| | 162 | 176 | HLA-DRB1*09:01 | NIAGLNALRLINDNT | 62 | 4.17 |
| | 158 | 172 | HLA-DRB1*13:02 | LDAANIAGLNALRLI | 99.3 | 6.04 |
| | 159 | 173 | HLA-DRB5*01:01 | DAANIAGLNALRLIN | 63.2 | 11.64 |
| | 158 | 172 | HLA-DRB5*01:01 | LDAANIAGLNALRLI | 73 | 12.73 |
| | 160 | 174 | HLA-DRB5*01:01 | AANIAGLNALRLIND | 75 | 12.95 |
| VVFGTANPI | 416 | 430 | HLA-DQA1*05:01/DQB1*03:01 | EDTELVVFGTANPIP | 31.4 | 5.95 |
| | 415 | 429 | HLA-DQA1*05:01/DQB1*03:01 | DEDTELVVFGTANPI | 39.1 | 7.29 |
| | 418 | 432 | HLA-DRB1*01:01 | TELVVFGTANPIPST | 9.5 | 4.8 |
| | 419 | 433 | HLA-DRB1*01:01 | ELVVFGTANPIPSTK | 12.4 | 6.94 |
| | 420 | 434 | HLA-DRB1*01:01 | LVVFGTANPIPSTKV | 13.4 | 7.6 |

| | | | | | | |
|---|---|---|---|---|---|---|
| | 417 | 431 | HLA-DRB1*01:01 | DTELVVFGTANPIPS | 15.9 | 9.13 |
| | 415 | 429 | HLA-DRB1*01:01 | DEDTELVVFGTANPI | 26.6 | 14.19 |
| | 416 | 430 | HLA-DRB1*01:01 | EDTELVVFGTANPIP | 28.8 | 15.03 |
| | 421 | 435 | HLA-DRB1*01:01 | VVFGTANPIPSTKVL | 34.1 | 16.85 |
| | 415 | 429 | HLA-DRB1*07:01 | DEDTELVVFGTANPI | 17.5 | 3.31 |
| | 416 | 430 | HLA-DRB1*07:01 | EDTELVVFGTANPIP | 20.7 | 3.99 |
| | 417 | 431 | HLA-DRB1*07:01 | DTELVVFGTANPIPS | 23 | 4.44 |
| | 418 | 432 | HLA-DRB1*07:01 | TELVVFGTANPIPST | 30.2 | 5.73 |
| | 420 | 434 | HLA-DRB1*07:01 | LVVFGTANPIPSTKV | 41.3 | 7.41 |
| | 419 | 433 | HLA-DRB1*07:01 | ELVVFGTANPIPSTK | 42.1 | 7.52 |
| | 421 | 435 | HLA-DRB1*07:01 | VVFGTANPIPSTKVL | 43.7 | 7.78 |
| | 418 | 432 | HLA-DRB1*13:02 | TELVVFGTANPIPST | 3.3 | 0.05 |
| | 417 | 431 | HLA-DRB1*13:02 | DTELVVFGTANPIPS | 3.5 | 0.06 |
| | 419 | 433 | HLA-DRB1*13:02 | ELVVFGTANPIPSTK | 3.7 | 0.07 |
| | 416 | 430 | HLA-DRB1*13:02 | EDTELVVFGTANPIP | 3.9 | 0.08 |
| | 415 | 429 | HLA-DRB1*13:02 | DEDTELVVFGTANPI | 4.3 | 0.11 |
| | 420 | 434 | HLA-DRB1*13:02 | LVVFGTANPIPSTKV | 4.4 | 0.12 |
| | 421 | 435 | HLA-DRB1*13:02 | VVFGTANPIPSTKVL | 5.5 | 0.2 |
| | 418 | 432 | HLA-DRB1*15:01 | TELVVFGTANPIPST | 71.1 | 7.28 |
| | 417 | 431 | HLA-DRB1*15:01 | DTELVVFGTANPIPS | 85 | 8.6 |
| | 419 | 433 | HLA-DRB1*15:01 | ELVVFGTANPIPSTK | 96.5 | 9.63 |
| **FYRQGAFEL*** | 433 | 447 | HLA-DPA1*01/DPB1*04:01 | KVLTFYRQGAFELEA | 67.8 | 3.94 |
| | 433 | 447 | HLA-DPA1*01:03/DPB1*02:01 | KVLTFYRQGAFELEA | 63.4 | 6.11 |
| | 432 | 446 | HLA-DPA1*01:03/DPB1*02:01 | TKVLTFYRQGAFELE | 69.2 | 6.5 |
| | 434 | 448 | HLA-DPA1*01:03/DPB1*02:01 | VLTFYRQGAFELEAA | 69.3 | 6.5 |
| | 435 | 449 | HLA-DPA1*01:03/DPB1*02:01 | LTFYRQGAFELEAAY | 91.2 | 7.86 |
| | 431 | 445 | HLA-DPA1*01:03/DPB1*02:01 | STKVLTFYRQGAFEL | 92.1 | 7.91 |
| | 434 | 448 | HLA-DPA1*02:01/DPB1*01:01 | VLTFYRQGAFELEAA | 36.5 | 3.66 |
| | 435 | 449 | HLA-DPA1*02:01/DPB1*01:01 | LTFYRQGAFELEAAY | 39.8 | 4.07 |
| | 433 | 447 | HLA-DPA1*02:01/DPB1*01:01 | KVLTFYRQGAFELEA | 41.3 | 4.25 |
| | 432 | 446 | HLA-DPA1*02:01/DPB1*01:01 | TKVLTFYRQGAFELE | 44.7 | 4.66 |
| | 436 | 450 | HLA-DPA1*02:01/DPB1*01:01 | TFYRQGAFELEAAYA | 72.9 | 7.82 |
| | 431 | 445 | HLA-DPA1*02:01/DPB1*01:01 | STKVLTFYRQGAFEL | 85.8 | 9.09 |
| | 434 | 448 | HLA-DRB1*01:01 | VLTFYRQGAFELEAA | 5.7 | 1.43 |
| | 433 | 447 | HLA-DRB1*01:01 | KVLTFYRQGAFELEA | 6.6 | 2.28 |
| | 435 | 449 | HLA-DRB1*01:01 | LTFYRQGAFELEAAY | 7.1 | 2.73 |
| | 432 | 446 | HLA-DRB1*01:01 | TKVLTFYRQGAFELE | 8.1 | 3.63 |
| | 431 | 445 | HLA-DRB1*01:01 | STKVLTFYRQGAFEL | 8.9 | 4.3 |
| | 436 | 450 | HLA-DRB1*01:01 | TFYRQGAFELEAAYA | 10.6 | 5.65 |
| | 437 | 451 | HLA-DRB1*01:01 | FYRQGAFELEAAYAD | 16.2 | 9.3 |
| | 431 | 445 | HLA-DRB1*07:01 | STKVLTFYRQGAFEL | 19.1 | 3.63 |
| | 432 | 446 | HLA-DRB1*07:01 | TKVLTFYRQGAFELE | 28.1 | 5.38 |
| | 433 | 447 | HLA-DRB1*07:01 | KVLTFYRQGAFELEA | 38.9 | 7.04 |
| | 434 | 448 | HLA-DRB1*07:01 | VLTFYRQGAFELEAA | 58.1 | 9.72 |

| | | | | | | |
|---|---|---|---|---|---|---|
| | 435 | 449 | HLA-DRB1*07:01 | LTFYRQGAFELEAAY | 94.1 | 13.61 |
| | 434 | 448 | HLA-DRB1*09:01 | VLTFYRQGAFELEAA | 51.5 | 3.35 |
| | 433 | 447 | HLA-DRB1*09:01 | KVLTFYRQGAFELEA | 58 | 3.86 |
| | 435 | 449 | HLA-DRB1*09:01 | LTFYRQGAFELEAAY | 70.3 | 4.79 |
| | 432 | 446 | HLA-DRB1*09:01 | TKVLTFYRQGAFELE | 71 | 4.83 |
| | 436 | 450 | HLA-DRB1*09:01 | TFYRQGAFELEAAYA | 98.9 | 6.84 |
| | 431 | 445 | HLA-DRB1*09:01 | STKVLTFYRQGAFEL | 99.1 | 6.86 |
| | 431 | 445 | HLA-DRB5*01:01 | STKVLTFYRQGAFEL | 63 | 11.62 |
| | 434 | 448 | HLA-DRB5*01:01 | VLTFYRQGAFELEAA | 79.2 | 13.37 |
| | 433 | 447 | HLA-DRB5*01:01 | KVLTFYRQGAFELEA | 84.4 | 13.87 |
| | 432 | 446 | HLA-DRB5*01:01 | TKVLTFYRQGAFELE | 96.5 | 14.99 |
| LREALNTYL | 655 | 669 | HLA-DRB1*01:01 | AAALREALNTYLTAA | 33.8 | 16.76 |
| | 656 | 670 | HLA-DRB1*01:01 | AALREALNTYLTAAQ | 40 | 18.64 |
| | 657 | 671 | HLA-DRB1*01:01 | ALREALNTYLTAAQG | 50.3 | 21.32 |
| | 654 | 668 | HLA-DRB1*01:01 | AAAALREALNTYLTA | 54.2 | 22.23 |
| | 658 | 672 | HLA-DRB1*01:01 | LREALNTYLTAAQGE | 71.3 | 25.66 |
| | 653 | 667 | HLA-DRB1*01:01 | RAAAALREALNTYLT | 84.4 | 27.88 |
| | 654 | 668 | HLA-DRB1*04:05 | AAAALREALNTYLTA | 44.9 | 4.28 |
| | 655 | 669 | HLA-DRB1*04:05 | AAALREALNTYLTAA | 47.2 | 4.54 |
| | 653 | 667 | HLA-DRB1*04:05 | RAAAALREALNTYLT | 47.8 | 4.61 |
| | 656 | 670 | HLA-DRB1*04:05 | AALREALNTYLTAAQ | 58.1 | 5.73 |
| | 652 | 666 | HLA-DRB1*04:05 | PRAAAALREALNTYL | 61.5 | 6.06 |
| | 657 | 671 | HLA-DRB1*04:05 | ALREALNTYLTAAQG | 80 | 7.86 |
| | 653 | 667 | HLA-DRB1*07:01 | RAAAALREALNTYLT | 32.6 | 6.14 |
| | 654 | 668 | HLA-DRB1*07:01 | AAAALREALNTYLTA | 32.9 | 6.18 |
| | 652 | 666 | HLA-DRB1*07:01 | PRAAAALREALNTYL | 34 | 6.35 |
| | 655 | 669 | HLA-DRB1*07:01 | AAALREALNTYLTAA | 36.2 | 6.65 |
| | 656 | 670 | HLA-DRB1*07:01 | AALREALNTYLTAAQ | 44.1 | 7.83 |
| | 657 | 671 | HLA-DRB1*07:01 | ALREALNTYLTAAQG | 55.7 | 9.44 |
| | 658 | 672 | HLA-DRB1*07:01 | LREALNTYLTAAQGE | 68.7 | 11.01 |
| | 655 | 669 | HLA-DRB1*13:02 | AAALREALNTYLTAA | 70.1 | 4.63 |
| | 654 | 668 | HLA-DRB1*13:02 | AAAALREALNTYLTA | 84.6 | 5.39 |
| | 656 | 670 | HLA-DRB1*13:02 | AALREALNTYLTAAQ | 97.8 | 5.97 |
| | 655 | 669 | HLA-DRB5*01:01 | AAALREALNTYLTAA | 39.6 | 8.49 |
| | 656 | 670 | HLA-DRB5*01:01 | AALREALNTYLTAAQ | 65.2 | 11.87 |
| | 654 | 668 | HLA-DRB5*01:01 | AAAALREALNTYLTA | 66.6 | 12.02 |
| FKNTVGSLK | 58 | 72 | HLA-DRB1*01:01 | TSNFKNTVGSLKRLI | 25.2 | 13.63 |
| | 59 | 73 | HLA-DRB1*01:01 | SNFKNTVGSLKRLIG | 46.2 | 20.3 |
| | 57 | 71 | HLA-DRB1*01:01 | ETSNFKNTVGSLKRL | 46.4 | 20.35 |
| | 60 | 74 | HLA-DRB1*01:01 | NFKNTVGSLKRLIGR | 82.8 | 27.61 |
| | 58 | 72 | HLA-DRB1*04:05 | TSNFKNTVGSLKRLI | 97.2 | 9.39 |
| | 58 | 72 | HLA-DRB1*07:01 | TSNFKNTVGSLKRLI | 16.2 | 3.04 |
| | 59 | 73 | HLA-DRB1*07:01 | SNFKNTVGSLKRLIG | 19.5 | 3.71 |
| | 57 | 71 | HLA-DRB1*07:01 | ETSNFKNTVGSLKRL | 20.1 | 3.86 |

| | | | | | | |
|---|---|---|---|---|---|---|
| | 60 | 74 | HLA-DRB1*07:01 | NFKNTVGSLKRLIGR | 23.9 | 4.61 |
| | 56 | 70 | HLA-DRB1*07:01 | AETSNFKNTVGSLKR | 29.3 | 5.58 |
| | 61 | 75 | HLA-DRB1*07:01 | FKNTVGSLKRLIGRS | 29.6 | 5.63 |
| | 55 | 69 | HLA-DRB1*07:01 | TAETSNFKNTVGSLK | 31.7 | 5.99 |
| | 58 | 72 | HLA-DRB1*09:01 | TSNFKNTVGSLKRLI | 76.7 | 5.28 |
| | 57 | 71 | HLA-DRB1*09:01 | ETSNFKNTVGSLKRL | 98.1 | 6.78 |
| | 58 | 72 | HLA-DRB5*01:01 | TSNFKNTVGSLKRLI | 4.1 | 0.57 |
| | 59 | 73 | HLA-DRB5*01:01 | SNFKNTVGSLKRLIG | 4.8 | 0.78 |
| | 57 | 71 | HLA-DRB5*01:01 | ETSNFKNTVGSLKRL | 5.3 | 0.93 |
| | 60 | 74 | HLA-DRB5*01:01 | NFKNTVGSLKRLIGR | 6.8 | 1.41 |
| | 56 | 70 | HLA-DRB5*01:01 | AETSNFKNTVGSLKR | 8 | 1.78 |
| | 61 | 75 | HLA-DRB5*01:01 | FKNTVGSLKRLIGRS | 10.1 | 2.4 |
| | 55 | 69 | HLA-DRB5*01:01 | TAETSNFKNTVGSLK | 16.4 | 4.05 |
| IVKVKARLN | 478 | 492 | HLA-DRB1*01:01 | DLSIVKVKARLNLHG | 14.2 | 8.11 |
| | 479 | 493 | HLA-DRB1*01:01 | LSIVKVKARLNLHGI | 16.6 | 9.52 |
| | 477 | 491 | HLA-DRB1*01:01 | GDLSIVKVKARLNLH | 16.8 | 9.63 |
| | 476 | 490 | HLA-DRB1*01:01 | SGDLSIVKVKARLNL | 20.9 | 11.74 |
| | 480 | 494 | HLA-DRB1*01:01 | SIVKVKARLNLHGIM | 25 | 13.55 |
| | 481 | 495 | HLA-DRB1*01:01 | IVKVKARLNLHGIMN | 32.3 | 16.25 |
| | 475 | 489 | HLA-DRB1*01:01 | ASGDLSIVKVKARLN | 33.1 | 16.52 |
| | 476 | 490 | HLA-DRB1*07:01 | SGDLSIVKVKARLNL | 8.7 | 1.34 |
| | 477 | 491 | HLA-DRB1*07:01 | GDLSIVKVKARLNLH | 9.9 | 1.62 |
| | 475 | 489 | HLA-DRB1*07:01 | ASGDLSIVKVKARLN | 10.3 | 1.72 |
| | 478 | 492 | HLA-DRB1*07:01 | DLSIVKVKARLNLHG | 13.5 | 2.46 |
| | 479 | 493 | HLA-DRB1*07:01 | LSIVKVKARLNLHGI | 14.4 | 2.67 |
| | 480 | 494 | HLA-DRB1*07:01 | SIVKVKARLNLHGIM | 18.3 | 3.46 |
| | 481 | 495 | HLA-DRB1*07:01 | IVKVKARLNLHGIMN | 22.4 | 4.33 |
| | 480 | 494 | HLA-DRB1*08:02 | SIVKVKARLNLHGIM | 57.3 | 0.62 |
| | 479 | 493 | HLA-DRB1*08:02 | LSIVKVKARLNLHGI | 65.9 | 0.77 |
| | 478 | 492 | HLA-DRB1*08:02 | DLSIVKVKARLNLHG | 78.5 | 1.06 |
| | 478 | 492 | HLA-DRB1*11:01 | DLSIVKVKARLNLHG | 11.4 | 1.65 |
| | 477 | 491 | HLA-DRB1*11:01 | GDLSIVKVKARLNLH | 12.7 | 1.95 |
| | 476 | 490 | HLA-DRB1*11:01 | SGDLSIVKVKARLNL | 13.8 | 2.22 |
| | 479 | 493 | HLA-DRB1*11:01 | LSIVKVKARLNLHGI | 14.1 | 2.28 |
| | 480 | 494 | HLA-DRB1*11:01 | SIVKVKARLNLHGIM | 20.7 | 3.73 |
| | 481 | 495 | HLA-DRB1*11:01 | IVKVKARLNLHGIMN | 30 | 5.51 |
| | 477 | 491 | HLA-DRB5*01:01 | GDLSIVKVKARLNLH | 35.4 | 7.82 |
| | 478 | 492 | HLA-DRB5*01:01 | DLSIVKVKARLNLHG | 36.2 | 7.95 |
| | 479 | 493 | HLA-DRB5*01:01 | LSIVKVKARLNLHGI | 39.8 | 8.52 |
| | 476 | 490 | HLA-DRB5*01:01 | SGDLSIVKVKARLNL | 44 | 9.15 |
| | 480 | 494 | HLA-DRB5*01:01 | SIVKVKARLNLHGIM | 61.3 | 11.41 |
| | 475 | 489 | HLA-DRB5*01:01 | ASGDLSIVKVKARLN | 68.7 | 12.27 |
| | 481 | 495 | HLA-DRB5*01:01 | IVKVKARLNLHGIMN | 95.9 | 14.94 |
| LKRLIGRSF | 63 | 77 | HLA-DRB1*01:01 | NTVGSLKRLIGRSFN | 18.2 | 10.39 |

| | | | | | |
|---|---|---|---|---|---|
| 64 | 78 | HLA-DRB1*01:01 | TVGSLKRLIGRSFND | 18.6 | 10.59 |
| 65 | 79 | HLA-DRB1*01:01 | VGSLKRLIGRSFNDP | 21 | 11.79 |
| 62 | 76 | HLA-DRB1*01:01 | KNTVGSLKRLIGRSF | 28.2 | 14.8 |
| 66 | 80 | HLA-DRB1*01:01 | GSLKRLIGRSFNDPE | 42.7 | 19.39 |
| 67 | 81 | HLA-DRB1*01:01 | SLKRLIGRSFNDPEV | 84.8 | 27.94 |
| 62 | 76 | HLA-DRB1*07:01 | KNTVGSLKRLIGRSF | 34.8 | 6.46 |
| 63 | 77 | HLA-DRB1*07:01 | NTVGSLKRLIGRSFN | 51.6 | 8.94 |
| 64 | 78 | HLA-DRB1*07:01 | TVGSLKRLIGRSFND | 70.2 | 11.18 |
| 65 | 79 | HLA-DRB1*09:01 | VGSLKRLIGRSFNDP | 32.4 | 1.76 |
| 64 | 78 | HLA-DRB1*09:01 | TVGSLKRLIGRSFND | 37 | 2.15 |
| 66 | 80 | HLA-DRB1*09:01 | GSLKRLIGRSFNDPE | 39.7 | 2.39 |
| 63 | 77 | HLA-DRB1*09:01 | NTVGSLKRLIGRSFN | 51.4 | 3.34 |
| 67 | 81 | HLA-DRB1*09:01 | SLKRLIGRSFNDPEV | 61.4 | 4.13 |
| 62 | 76 | HLA-DRB1*09:01 | KNTVGSLKRLIGRSF | 71.6 | 4.88 |
| 65 | 79 | HLA-DRB1*11:01 | VGSLKRLIGRSFNDP | 8.9 | 1.06 |
| 64 | 78 | HLA-DRB1*11:01 | TVGSLKRLIGRSFND | 10.8 | 1.51 |
| 66 | 80 | HLA-DRB1*11:01 | GSLKRLIGRSFNDPE | 11.8 | 1.74 |
| 67 | 81 | HLA-DRB1*11:01 | SLKRLIGRSFNDPEV | 16.7 | 2.85 |
| 68 | 82 | HLA-DRB1*11:01 | LKRLIGRSFNDPEVE | 26.6 | 4.9 |
| 62 | 76 | HLA-DRB5*01:01 | KNTVGSLKRLIGRSF | 42.5 | 8.94 |
| 64 | 78 | HLA-DRB5*01:01 | TVGSLKRLIGRSFND | 51.2 | 10.15 |
| 63 | 77 | HLA-DRB5*01:01 | NTVGSLKRLIGRSFN | 52.1 | 10.27 |
| 65 | 79 | HLA-DRB5*01:01 | VGSLKRLIGRSFNDP | 68.5 | 12.25 |

*Top promising epitopes with efficient binding affinity
