## Supplementary material for "Computational vaccinology approach: Designing an efficient multi-epitope peptide vaccine against *Cryptococcus neoformans var. grubii’s* heat shock 70KDa protein": supplemantal Table 6

### Supplementary Tables and Figures

**Table 6.** List of the promising epitopes core sequence that had an efficient binding affinity with MHC-I & MHC-II in combined mode.

| Core Sequence | Allele |
|---|---|
| **FYRQGAFEL*** | HLA-A*23:01, HLA-A*24:02, HLA-C*03:03, HLA-C*07:02, HLA-C*12:03, HLA-C*14:02,HLA-DPA1*01, HLA-DPB1*04:01, HLA-DPA1*01:03, HLA-DPB1*02:01, HLA-DPA1*02:01,HLA-DPB1*01:01, HLA-DRB1*01:01, HLA-DRB1*07:01, HLA-DRB1*09:01, HLA-DRB5*01:01 |
| FTQLVAAYL | HLA-A*02:01, HLA-A*02:06, HLA-A*68:02, HLA-C*03:03, HLA-C*05:01, HLA-C*14:02, HLA-C*15:02, HLA-DRB1*01:01, HLA-DRB1*04:04, HLA-DRB1*04:05, HLA-DRB1*07:01, HLA-DRB1*09:01, HLA-DRB1*11:01, HLA-DRB1*15:01, HLA-DRB5*01:01 |
| FDYALVQHF | HLA-B*15:01, HLA-C*12:03, HLA-C*14:02, HLA-DPA1*01:03, HLA-DPB1*02:01, HLA-DPA1*02:01, HLA-DPB1*01:01, HLA-DRB1*01:01, HLA-DRB1*03:01, HLA-DRB1*04:05, HLA-DRB1*07:01, HLA-DRB1*09:01, HLA-DRB1*11:01, HLA-DRB5*01:01 |
| **FFGGKVLNF*** | HLA-A*23:01, HLA-A*29:02, HLA-DPA1*01:03, HLA-DPB1*02:01, HLA-DPA1*02:01, HLA-DPB1*01:01, HLA-DPA1*03:01, HLA-DPB1*04:02, HLA-DQA1*05:01, HLA-DQB1*03:01, HLA-DRB1*01:01,HLA-DRB1*07:01 |
| YVYDTRGKL | HLA-A*02:06, HLA-A*68:02, HLA-B*07:02, HLA-C*03:03, HLA-C*06:02, HLA-C*07:01, HLA-C*12:03, HLA-C*14:02, HLA-C*15:02, HLA-DRB1*03:01, HLA-DRB1*11:01, HLA-DRB3*01:01 |
| FACASLSPV | HLA-A*02:01, HLA-A*02:06, HLA-A*68:02, HLA-C*03:03, HLA-C*12:03, HLA-C*15:02, HLA-DQA1*05:01, HLA-DQB1*03:01, HLA-DRB1*01:01, HLA-DRB1*07:01, HLA-DRB1*09:01 |
| **FINAQLVDV*** | HLA-A*02:01, HLA-A*02:06, HLA-A*68:02, HLA-DPA1*01:03, HLA-DPB1*02:01, HLA-DPA1*02:01, HLA-DPB1*01:01, HLA-DPA1*03:01, HLA-DPB1*04:02, HLA-DRB1*01:01 |
| LVQHFAEEF | HLA-B*35:01, HLA-DPA1*01:03, HLA-DPB1*02:01, HLA-DPA1*02:01, HLA-DPB1*01:01, HLA-DQA1*05:01, HLA-DQB1*02:01, HLA-DRB1*09:01 |
| FSFTQLVAA | HLA-A*02:01, HLA-A*02:06, HLA-A*68:02, HLA-DPA1*01:03, HLA-DPB1*02:01, HLA-DQA1*01:02, HLA-DQB1*06:02, HLA-DRB1*01:01 |

*Top promising epitopes with efficient binding affinity and massive population coverage
