## Supplementary material for "Computational vaccinology approach: Designing an efficient multi-epitope peptide vaccine against *Cryptococcus neoformans var. grubii’s* heat shock 70KDa protein": supplemantal Figure 1

### Supplementary Tables and Figures

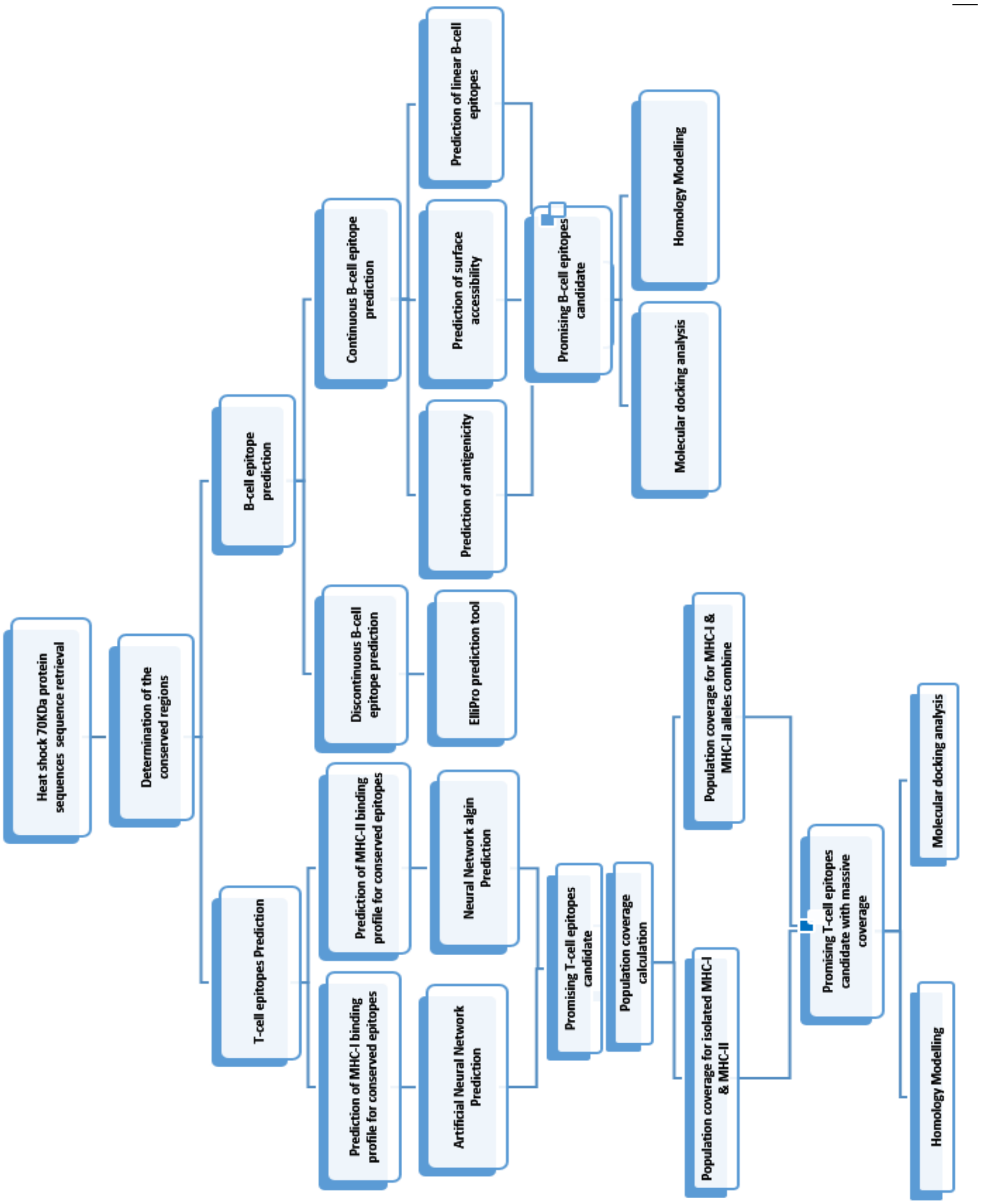


**Figure 1.** Graphical representation of Peptide vaccine designing against *C. neoformans's* heat shock 70KDa protein
